# Sorafenib induces muscle wasting by disrupting the activity of distinct chromatin regulators

**DOI:** 10.1101/2024.01.04.574149

**Authors:** Bushra Khan, Chiara Lanzuolo, Valentina Rosti, Philina Santarelli, Andreas Pich, Theresia Kraft, Mamta Amrute-Nayak, Arnab Nayak

## Abstract

Adverse effects of chemotherapies can outweigh the benefits in cancer patients. Various chemotherapeutics are linked to muscle wasting or cachexia, drastically reducing the chance of survivability of cancer patients. Insights into the molecular basis of chemotherapy-induced cachexia is an unmet need to improve the treatment strategies. Here, we investigated the tyrosine kinase inhibitor class of chemotherapeutic agents for their effects on muscle function. Sorafenib, but not Nilotinib and Imatinib, triggered cachexia. System-wide transcriptome and proteome analyses revealed that Sorafenib alters the global transcriptional program and proteostasis in muscle cells. Mechanistically, Sorafenib treatment reduced active epigenetic mark H3K4 methylation on distinct muscle-specific genes due to the defective chromatin association of SET1/A, a catalytic component of the SET1/MLL complex. It favored transcriptionally incompetent chromatin, characterized by diminished association with RNA polymerase II. The transcriptional reorientation led to disrupted sarcomere organization, calcium homeostasis, and mitochondrial respiration. Consequently, the contractile ability of muscle cells was severely compromised. Collectively, we identified an unanticipated transcriptional mechanism underlying Sorafenib-induced cachexia. Our findings hold the potential to strategize therapy regimens to minimize chemotherapy-induced cachexia and improve treatment outcomes.

## Introduction

Movement is among the defining metazoan features critically associated with their existence and evolution. A sophisticated and highly efficient form of movement is powered by skeletal muscle, which constitutes approximately 40% of total body weight and 50-70% of all proteins in humans ^1,2^. ATP-dependent actomyosin cross-bridge cycling within sarcomere, the fundamental contractile unit of skeletal muscle, produces force to bear load and drive movement ^3,4^. A precise organization of the sarcomere is key for optimal force generation and muscle contraction. In addition to an accurate sarcomere organization, sarcoplasmic reticulum (SR)-mediated calcium handling and mitochondrial energy production facilitate the contractile ability of muscle cells. Thus, a functional homeostasis, primarily between these organelles, is essential for muscle contraction. Deregulation of this interplay can lead to defective muscle function and muscle-wasting disorders such as cachexia ^5–7^. Cachexia is a multifactorial muscle wasting disorder, marked by severe involuntary loss of skeletal muscle mass in affected individuals, by approximately 30% of their pre-illness level ^8^. Cachexia is observed in ∼80% of advanced cancer patients and is the cause of mortality in more than 30% patients ^9^. By secreting a multitude of pro-inflammatory cytokines *viz*, tumor-necrosis factor (TNF-α), interleukin (IL)-1 etc., into their micro-environment, cancers can induce muscle protein catabolism in a p38 MAPK (mitogen-activated protein kinase)- and nuclear factor ‘kappa-light-chain-enhancer’ of activated B-cells (NF-κB)-dependent manner. This alters muscle metabolism, triggers degradation of muscle proteins and thereby induces cachexia, also termed as cancer-induced cachexia ^10–16^. In addition to cancer, the chemotherapeutic drugs used in cancer therapy have been recently reported as inducers of cachexia ^5,17–19^. This greatly exacerbates the already deteriorated situation and contributes to poor survival rate. Currently, there are no effective medical strategies to ameliorate cachexia, suggesting the need to improve our understanding of the mechanisms underlying cachexia. Earlier reports showed that chemotherapeutic agent daunorubicin triggered cachexia by increasing oxidative stress and activation of NFκB/MuRF1-dependent ubiquitin–proteasome pathway that stimulates protein catabolism ^20,21^. Interestingly, another study showed that cisplatin-induced cachexia is NFκB dependent-but independent of MuRF1 ^22^. The ambiguous reports led us to hypothesize that other unknown molecular pathways may be involved in causing cachexia and that different chemotherapy agents may exert differential effects on the corresponding pathways.

In this study, we investigated the tyrosine kinase inhibitor (TKI) family of chemotherapeutic drugs including Sorafenib (Sor), Imatinib (Ima) and Nilotinib (Nilo). TKIs constitute a repertoire of chemotherapeutic drugs that have shown great promise in treating diverse human cancers ^23^. They inhibit various kinase families, including vascular endothelial growth factor receptor (VEGFR), platelet-derived growth factor receptor (PDGFR) kinases and B/C-Raf kinases ^24^. TKI’s involvement in normal muscle function and in pathological conditions, however, remained poorly understood. Cascade of tightly controlled epigenetic mechanisms mediated by Trithorax (TrxG) and Polycomb (PcG) group of proteins ensures expression of homeobox (*HOX)* genes ^25–30^ essential for regulation of early stages of metazoan development. SET1/MLL (mixed-lineage leukemia histone methyltransferase complex) belongs to the TrxG group of epigenetics modifiers, and consists of six different protein isoforms. The core catalytic subunit is made up of one of the MLL isoforms (MLL1-4) or the SET1 isoforms (SET1A-B).The catalytic core is further associated with four subunits that form the WRAD module (WDR5, RbBP5, ASH2L, and DPY30), which are essential for H3K4me2 and H3K4me3 modifications ^31–35^. In addition to the WRAD complex, distinct MLL family members are associated with other regulatory components (such as Menin, WDR82 etc.) which can act as a scaffold for recruiting specific transcription factors ^32,36,37^. Regulation of target gene specificity of the SET1/MLL complex and the molecular mechanism of its chromatin association remains poorly determined. Moreover, the potential involvement of SET1/MLL complex in chemotherapy-induced cachexia is still unknown.

In this study, we demonstrated specific molecular mechanisms governed by Sor in regulating diverse aspects of muscle cell physiology and in cachexia. We identified that Sor alters the cellular transcriptional program by modulating SET1/MLL histone methyltransferase-governed epigenetic mechanisms on distinct muscle-specific genes that leads to reduced association of transcriptionally active RNA polymerase II on these genes, subsequently resulting in insufficient protein expression. Additionally, Sor triggers an imbalance of proteostasis in muscle cells. At the cellular level, Sor impedes functionally linked physiological pathways of the SR and mitochondria, which ultimately impinge on contractile and metabolic dysfunction of muscle cells. In contrast to Sor, Nilo and Ima, showed no significant adverse alteration of muscle function.

## Results

### Effect of TKIs on sarcomere organization and contractile propensity of muscle cells

To investigate the effects of TKIs on muscle cell physiology and function, we used C2C12 mouse progenitor cells and differentiated to form mature myotubes (Fig. 1A). The myotubes were treated with TKIs for 24 h and DMSO was used as the vehicle control. The drug concentrations used in this study were in accordance with previously optimized concentrations for mammalian cells, including muscle cells ^38,39^. We stained the sarcomeric Z-disc protein, alpha-actinin in immunofluorescence assays to check if TKIs induce any phenotypic changes in myotubes. Similar to the control cells, Nilo- and Ima-treated cells displayed a long, tubular structure. Myotubes treated with Sor transformed into a short, thin and spindle-shaped structure (Fig. 1B). Compared to DMSO-treated cells; we observed a significant reduction in myotube length upon Sor treatment (DMSO = 151.2 µm, Sor = 60.21 µm) (Fig. 1C). Measurement of the myotube diameter confirmed significant thinning of the myotubes upon Sor treatment (DMSO = 10.03 µm, Sor = 4.93 µm) (Fig. 1D). However, the myotube morphology, length (mean values of DMSO = 151.2 µm, Ima = 179.3 µm, Nilo = 149.6 µm) and diameter (DMSO = 10.03 µm, Ima = 9.11 µm, Nilo = 9.45 µm) remained largely unaltered upon Nilo and Ima treatment (Fig. 1C, D). The changed morphology of muscle cells might have consequences on muscle cell function. To test this notion, we monitored the contractile ability of TKI-treated myotubes using the MyoPacer device. The cells were electrically paced at 40 V and 1 Hz to stimulate contractions. The electric stimulation mimics a nerve impulse, triggering Ca^2+^ release from SR into the sarcoplasm and driving actomyosin cross-bridge cycling and thereby sarcomere contraction. Representative examples of periodic sarcomere contraction corresponding to the stimulus frequency is presented in Fig. 1E. Kymographs (time vs. distance plot) generated from multiple movies per treatment condition showed horizontal peaks at regular intervals, indicative of regular contraction of the myotubes. Approximately 70% of DMSO-treated cells showed contractility, which was comparable to Nilo treatment (∼63% contractile cells). A moderate-but insignificant-decrease in contractile cell population was observed in Ima treatment (∼54%) (Fig. 1F and Movie S1-3). Strikingly, only 0.15 % myotubes (i.e., 1 out of 678 cells from three independent experiments) showed periodic contractions upon Sor treatment (Fig. 1F and Movie S1-3).

**Fig. 1:**
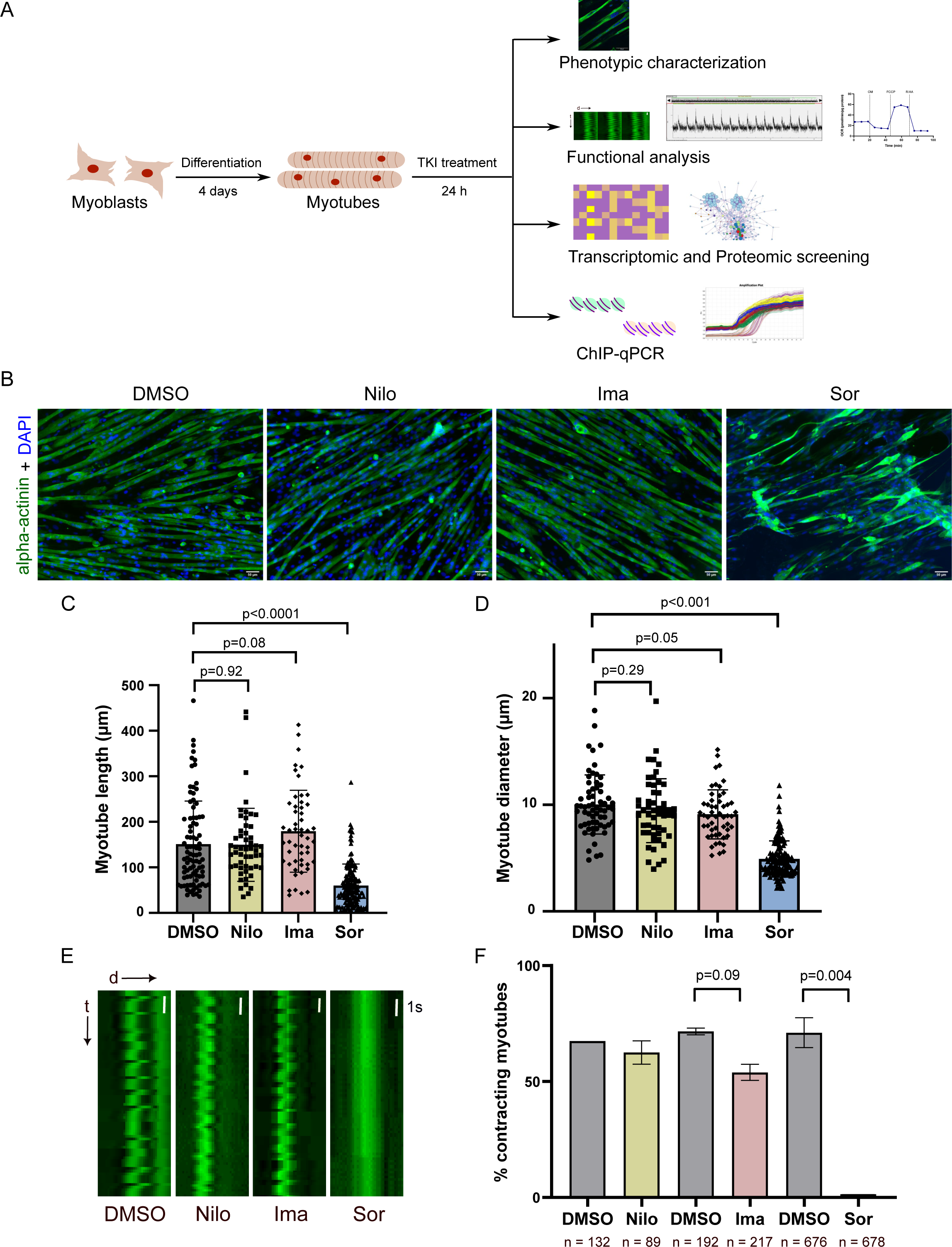
Effect of TKIs on cellular phenotype and function of myotubes. (A) Overview of the experimental plan used in this study. C2C12 mouse progenitor myoblast cells were differentiated for 4 days, followed by 24 h treatment with TKIs – Nilo, Ima and Sor. DMSO was used as the control. TKI-treated muscle cells were used for various assays as indicated (B) Representative images of TKI-treated myotubes immunostained against alpha-actinin (green) and nucleus (DAPI – blue). Images were captured using the 10X objective. Scale bar = 50 µm. (C) Graph shows myotube length (in µm) as calculated from images in (B). Note that the average myotube lengths presented in the plots for control, Ima, and Nilo are underestimated, as a large fraction of myotubes extend beyond even the lower magnified field of view and thus could not be measured. Therefore, the difference with the Sor-treated cells could be even larger. We monitored 89 cells for DMSO, 52 for Nilo, 52 cells for Ima, and, 150 cells for Sor treatment. (D) Graph shows myotube diameter (in µm) as calculated from images in (B). (E) Kymograph (time vs. distance plot) represents the contractility of myotubes upon electrical stimulation (generated from supplementary movies 1 - 3). The zigzag lines along the y-axis indicate periodic positional switches of specific structures, presumably organelles, along the length of the tubular cell. Horizontal intensity peaks for DMSO-, Nilo- and Ima-treated cells were observed at regular intervals. For Sor-treated cells, a straight line was observed, which indicates no change in the position of the corresponding structure at regular intervals, suggesting an inability of the cells to respond to stimuli and an absence of contraction-relaxation cycles. (F) Graph shows the percentage of contracting myotubes from pre-recorded movies at the MyoPacer, under different treatment conditions. Data is shown as Mean ± SEM, n - the total number of cells counted during the analysis. Statistical significance was determined using unpaired t-test with Welch’s correction on GraphPad Prism 9.5.1.

A precise sarcomere organization ensures proper contractile function of muscle cells. In order to understand the reasons underlying Sor-induced contractile defect, we stained TKI-treated myotubes with alpha-actinin, a Z disc protein, to visualize the sarcomere assembly ^40–42^. High-resolution confocal images showed a characteristic striated pattern indicating normal sarcomere organization in DMSO-treated control cells. Nilo and Ima treatment, which did not cause any contractile defect, also displayed well-organized sarcomere organization comparable to control cells. However, Sor-treated myotubes showed severely disordered sarcomere assembly (Fig. 2A and S1A). The defective sarcomere organization is unlikely to occur from changes in alpha-actinin protein stability since none of the three TKIs showed any measurable effect on the protein levels of alpha-actinin or myotilin, another prominent Z-disc protein (Fig. S1B-D). Collectively, these observations suggest that Sor-but not Nilo and Ima-significantly alters myotube morphology, disrupts sarcomere organization and, thereby, severely impairs contractile function of muscle cells.

**Fig. 2:**
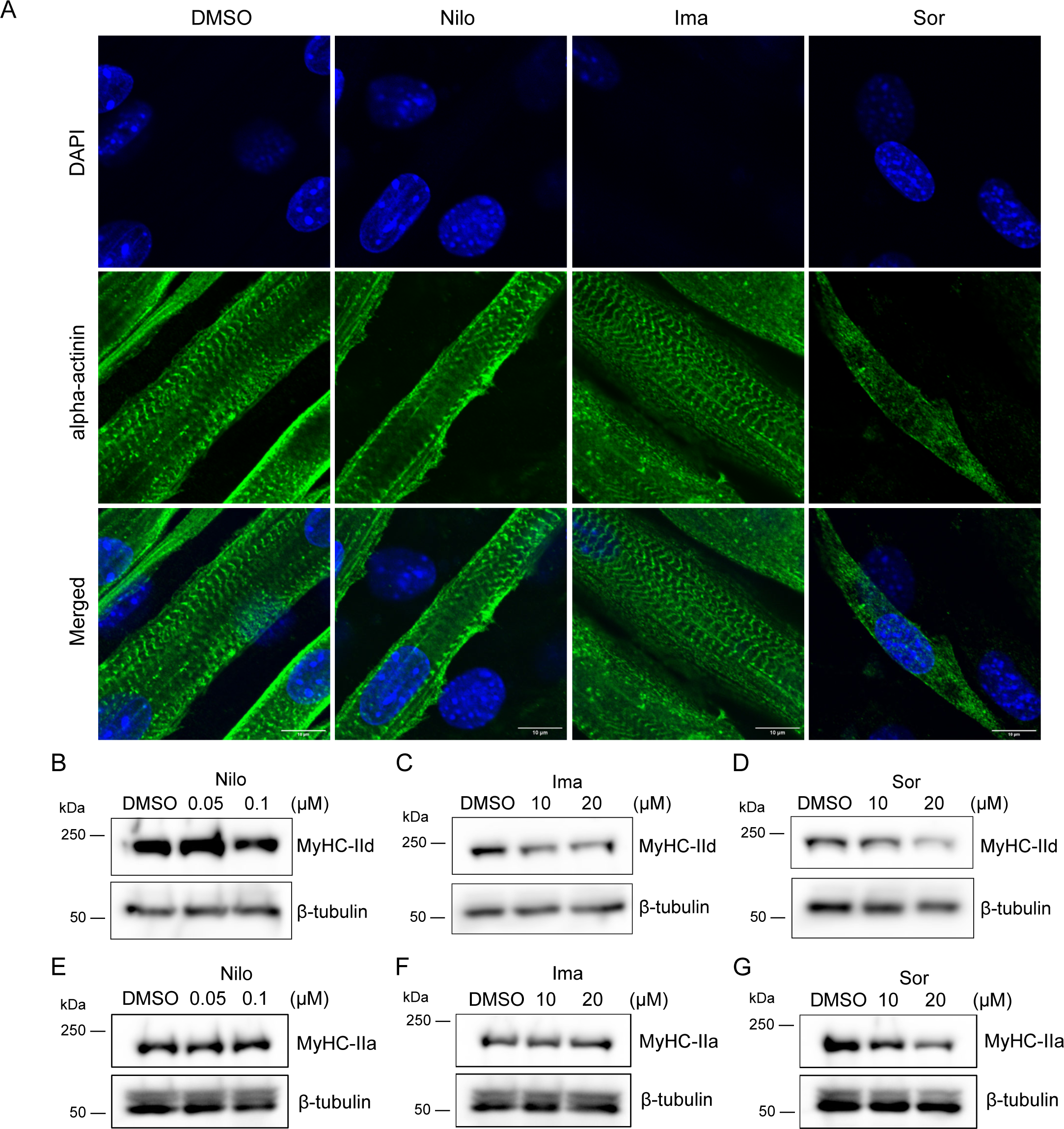
Effect of TKIs on sarcomere organization. (A) Panel shows confocal microscopy images of TKI-treated myotubes immunostained against alpha-actinin (green) and nucleus (DAPI – blue). Images were captured at 4X magnification over 60X oil immersion objective. Scale bar = 10 µm. See also Fig. S1. The images are the representative of at least three biological experiments. The concentrations used for Nilo was 0.1 µM, and 20 µM for Ima and Sor. (B–D) Western blot showing protein expression of MyHC-IId in myotubes treated with (B) Nilo, (C) Ima, and (D) Sor. (E-G) Western blot showing protein expression of MyHC-IIa in myotubes treated with (E) Nilo, (F) Ima, and (G) Sor. β-tubulin was used as the loading control. All concentrations are in µM.

### Sorafenib induces cachexia

Our observations of defective sarcomere organization and contractile dysfunction in Sor-treated muscle cells indicated Sor-triggered cachectic phenotype. Destabilization of myosin heavy chain II (MyHC II) protein has been reported as one of the hallmarks of cachexia ^43,44^. Thus, we sought to monitor the expression of the thick filament protein MyHC II in TKI-treated myotubes. Western blot analyses revealed that Sor treatment reduced the protein level of myosin isoforms MyHC-IId and MyHC-IIa, in a concentration-dependent manner (Fig. 2D, G). Note that the 20μM Sor concentration that we used in all our assays here is within the recommended effective dose in mammalian cells including muscle cells ^38,39^. The same was the case for Ima and Nilo doses. Interestingly, Nilo and Ima treatment did not alter MyHC-IIa level (Fig. 2 E, F). The effect of Nilo on MyHC-IId protein level was also not comparable to Sor (Fig. 2B). Ima showed only a mild effect MyHC-IId protein level. Collectively, these results indicate that specific TKIs i.e., Sor-but not Nilo and Ima-has major effect in reducing myosin heavy chain protein expression, which is a protagonist motor protein component, thereby, impairing myotube contractile ability. The mild effect of Ima was not significant enough to impair sarcomere organization and cell contraction. Since we detected a predominant cachectic effect of Sor, we focused on understanding the mechanisms underlying Sor-induced cachexia.

### Sorafenib treatment abrogates sarcoplasmic reticulum (SR)-mediated calcium handling

The contractile feature and force generation capacity of muscle cell is critically dependent on sarcomere organization and SR-mediated calcium handling. This entails Ca^2+^ release via Ryanodine receptor 1 (RyR1) and Ca^2+^ re-uptake via Sarco/endoplasmic reticulum calcium ATPase 1 (SERCA1), thereby creating a transient of Ca^2+^ concentration in the cytosol ^45,46^. Increased Ca^2+^ concentration in sarcoplasm critically activates the acto-myosin cross-bridge cycling, and thereby, sarcomere shortening and force generation by striated muscle cells. The reuptake of Ca^2+^ into the SR via SERCA pump results in sarcomere/muscle relaxation. To understand the implications of Sor treatment on this regulatory aspect of muscle cell contraction, we measured single cell Ca^2+^ transients in differentiated myotubes. Upon electric stimulation, 97 % control myotubes showed a detectable Ca^2+^ transient with a strong Ca^2+^ release (visualized as the peak height) and reuptake (fluorescence decay) that resulted in myotube contraction and relaxation, respectively (Fig. 3A and B). Strikingly, Sor-treated myotubes displayed no analysable calcium transients. Thus, Sor affected SR-governed Ca^2+^ handling and destabilized cellular calcium homeostasis.

**Fig. 3:**
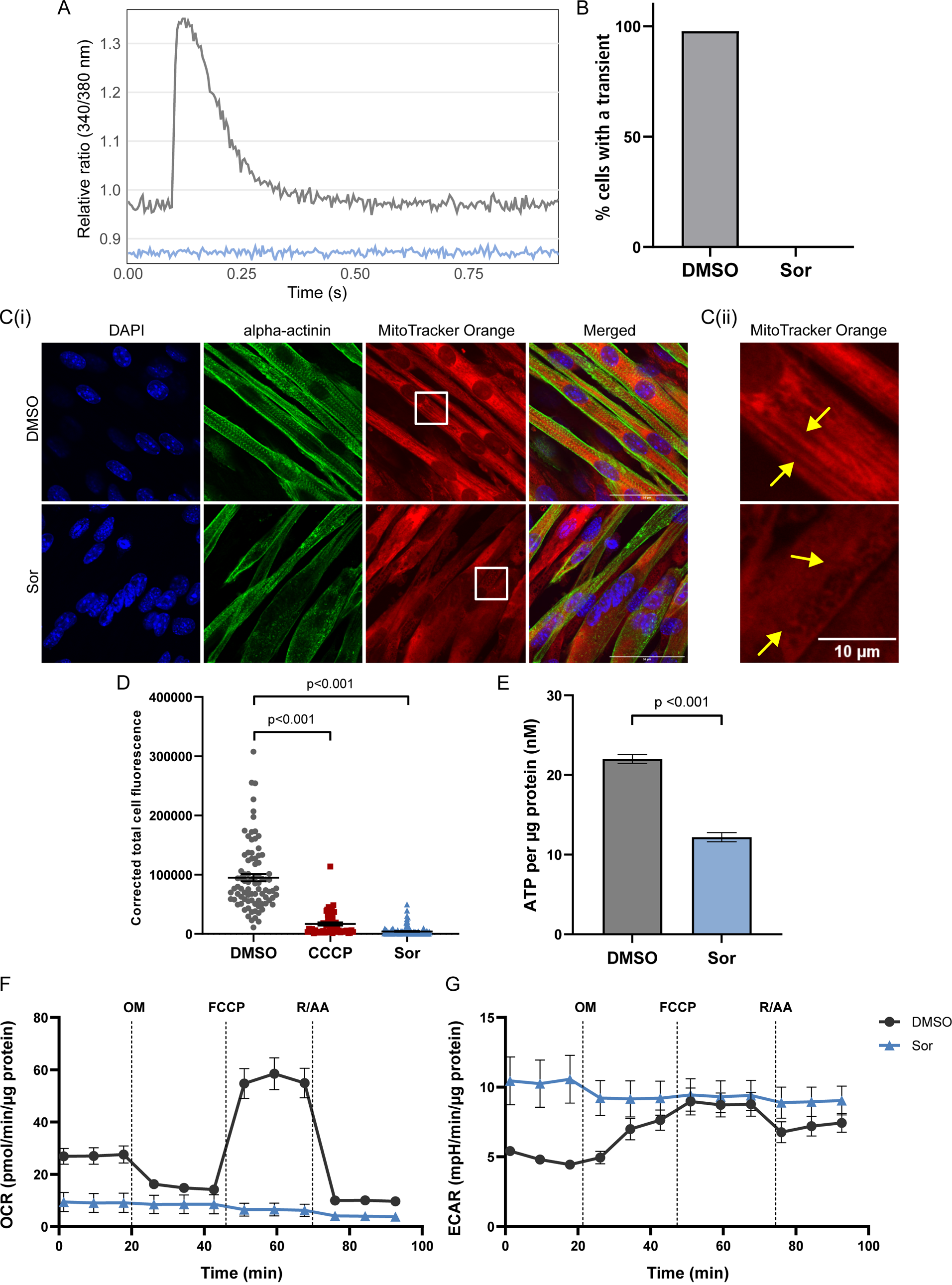
Sorafenib impairs Ca^2+^ homeostasis and mitochondrial function in muscle cells. (A) Representative single cell Ca^2+^ transient recorded over 90 sec, of DMSO- and Sor-treated muscle cell in grey and blue, respectively. (B) Bar graph showing the percentage of cells that showed a Ca^2+^ transient, where n = 44 for DMSO and n = 49 for Sor. The data were collected from two independent biological experiments. (C.i) Panel shows myotubes immunostained against alpha-actinin (green), nucleus (DAPI – blue) and mitochondria (MitoTracker Orange – red). Scale bar = 50 µm. (C.ii) Panel shows magnified white rectangular inset from the MioTracker channel in (C.i). Images were captured using the 60X oil immersion objective. Scale bar = 10 µm. (D) Scatter plot shows corrected total cell fluorescence of TMRM in myotubes treated with DMSO and Sor, as calculated from images in Fig. S2A. CCCP was used as the positive control of mitochondrial membrane depolarization. Data is represented as Mean ± SEM across three biological replicates, where n = 85 for DMSO, n = 56 for CCCP and n = 109 for Sor. (E) Graph shows quantification of total ATP per µg protein (in nM) in DMSO- and Sor-treated myotubes (Mean ± SEM, N = 3, n = 12, where N= biological replicates and n= technical replicates). (F) Graph shows time-course measurement of oxygen consumption rate (OCR), and (G) extracellular acidification rate (ECAR), in myotubes treated with DMSO and Sor (Mean ± SEM, N = 4, n = 44, where N= biological replicates and n= technical replicates). OM = oligomycin, FCCP = carbonyl cyanide-p-trifluoromethoxyphenylhydrazone, R/AA = rotenone/antimycin A. See also Fig. S2. Statistical significance was determined using unpaired t-test with Welch’s correction on GraphPad Prism 9.5.1. Control DMSO treatment= grey, Sor treatment = blue.

### Sorafenib induced cachexia entails mitochondrial dysfunction

Calcium homeostasis in muscle tissue is maintained by interplay between the SR and mitochondria ^47^. Since Sor impaired SR-dependent Ca^2+^ transient, we investigated the effects of Sor on mitochondrial function. As a first step, we monitored mitochondrial localization by MitoTracker Orange staining. Confocal images revealed diffused mitochondrial staining in Sor-treated myotubes, in contrast to the thread-like distribution that was observed in control cells (Fig. 3C). Noteworthy, alpha-actinin staining in this set up again confirmed our previous observations of disrupted sarcomere assembly upon Sor treatment. To further investigate Sor-triggered defects in mitochondrial function, we measured inner mitochondrial membrane integrity by staining the cells with the cationic dye TMRM (Tetramethylrhodamine methyl ester) and monitored its accumulation within the mitochondria. Typically, in healthy cells, the hyperpolarized inner mitochondrial membrane allows accumulation of TMRM within the mitochondrial matrix, resulting in higher TMRM fluorescence. In contrast, metabolically stressed cells display mitochondrial membrane potential (Δψ_m_) collapse and the dye is dispersed throughout the cytosol, resulting in lower fluorescence as shown with CCCP control (Fig. 3D). When compared to the control cells, quantification of total cell fluorescence revealed significantly lower TMRM accumulation (∼67.4 fold) in Sor-treated cells, suggesting alteration of the mitochondrial membrane potential (Fig. 3D, S2A). Since, changes in Δψ_m_ is critical for ATP production by the mitochondria, we checked whether Sor has any effect on ATP production. We observed a significant reduction in ATP production by Sor-treated muscle cells (Fig. 3E), indicative of perturbed mitochondrial function. As the ATP production is a direct result of oxidative phosphorylation (OXPHOS), we then checked mitochondrial respiratory capacity by employing the Seahorse Mito Stress test wherein the oxygen consumption rate (OCR) and extracellular acidification rate (ECAR) of the mitochondria is measured. Treatment of muscle cells with Sor significantly impaired the mitochondrial respiratory capacity (Fig. 3F) and reduced the basal respiration by ∼3 fold (Fig. S2B). Additionally, Sor-treated muscle cells showed very high ECAR values, indicating a metabolic switch to the energy-inefficient glycolytic mode of respiration (Fig. 3G). This was further validated by a reduction in coupling efficiency (an estimate of oxidative respiration used to drive ATP synthesis) in Sor-treated muscle cells (Fig. S2C), and a negative spare respiratory capacity (indicates the ability of a cell to meet metabolic requirements under stress conditions) (Fig. S2D). Taken together, these findings indicate that Sor induces cachexia by exerting multi-faceted effects on interconnected physiological pathways i.e., sarcomere organization, calcium handling, and mitochondrial metabolic activity.

### Sorafenib causes transcriptional reprogramming in muscle cells

To further examine the molecular mechanisms governing Sor-induced cachexia, we performed transcriptome profiling of muscle cells in response to Sor treatment. As a measure of data quality control, we computed the overall similarity between the biological replicates of the Sor-treated myotube populations by plotting a sample-to-sample distance heatmap. The heatmap showed strong clustering within the control and treatment samples, suggesting high degree of similarity and consistency among the different experiments (Fig. 4A). The distribution of gene expression in Sor-treated muscle cells as compared to the control was visualized using the MA plot, where the mean expression vs log_2_-fold change of each gene is plotted (Fig. 4B). The volcano plot showed 2128 genes as upregulated, while a substantially higher number of genes (3030) were downregulated (Fig. 4C). GO (Gene Ontology) enrichment analyses of the differentially regulated genes using DAVID software ^48^ revealed that most of the downregulated genes were predominantly clustered in GO terms including sarcomere organization and sarcoplasmic reticulum (Fig. 4D). Molecular motor protein-coding genes such as myosin heavy chain (*Myh1/MyHC-IId*, *Myh2/MyHC-IIa*) were among the most significantly downregulated genes. Interestingly, the myosin light chain genes such as myosin light chain 1 (*Myl1)* was also significantly downregulated. Moreover, genes encoding sarcomeric thin filament proteins (such as *Tnnt1*, *Acta1*, *Neb*, *Tnni1)*, M-band proteins (*Myom2*, and *Ttn)* were also downregulated (Table S1). The SR genes (such as *Ryr1, Atp2a1, Casq1 and several DHPR subunits*) associated with calcium transient processes represented another predominantly downregulated gene cluster in Sor-treated muscle cells (Fig. 4E, Table S1). Importantly, Sor treatment also led to the downregulation of protein-coding genes of the mitochondrial respiratory chain complex I, III, IV and V subunits (*mt-Nd1*, *mt-Nd6*, *Ndufs2*, *mt-Cytb*, *Cox7a1*, *mt-Atp6* among others) (Fig. 4F). On the contrary, the upregulated genes belonged to GO terms-endoplasmic reticulum (ER) (such as *Dnajc1*, *Tram1*, *Derl1*) and response to unfolded protein (including *Atf6*, *Herpud1*, *Hspa1b*, *Hspd1*, *Ddit3*) (Fig. 4C, D). In summary, Sor deregulates transcriptional processes of genes crucial for muscle contraction, calcium homeostasis, as well as mitochondrial oxidative respiration.

**Fig. 4:**
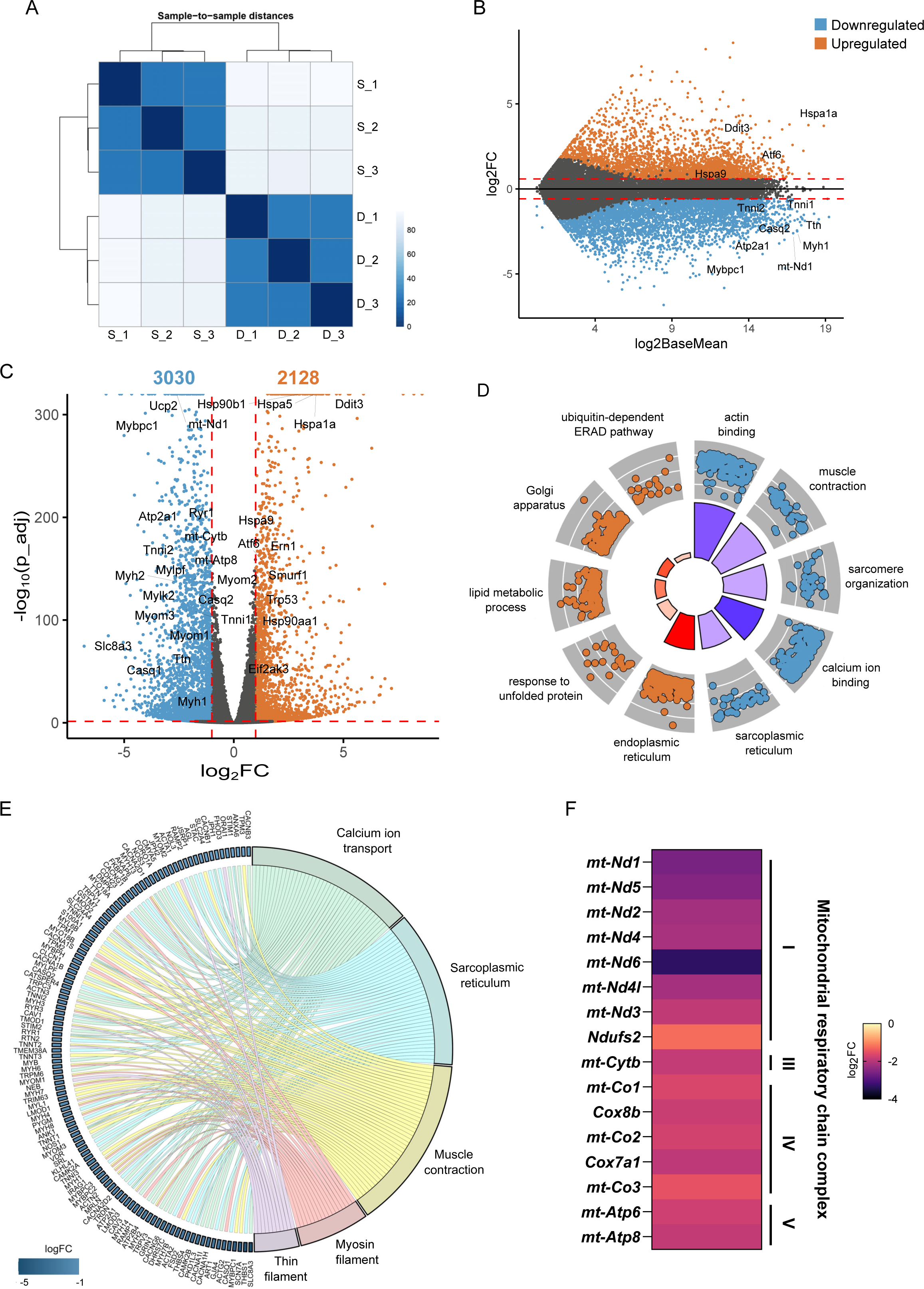
Transcriptomic profiling by RNAseq reveals deregulation of key cellular pathways in Sor-treated muscle cells. (A) Hierarchical clustering shows sample-to-sample distance between the different replicates, calculated based on the Euclidean distances between the DESeq2 rlog values of each sample. D_1, D_2 and D_3 represent three replicates of DMSO treatment, and S_1, S_2 and S_3 represent Sorafenib treatment. (B) MA plot of log_2_-fold change vs log_2_-base mean expression of all genes. Differentially regulated genes were identified by filtering the genes at a threshold p-value ≤ 0.05 and log2-fold change ≥ ±1.0, as indicated by the dotted lines. Each dot represents one gene, upregulated genes are shown in orange and downregulated genes are in blue. (C) Volcano plot shows the total number of up- and down-regulated genes in Sor-vs DMSO-treated myotubes. Dotted lines indicate the threshold fold change and p-value used (p-value ≤ 0.05 and log2-fold change ≥ ±1.0) for filtering the dataset. Each dot represents one gene. (D) Circle plot shows the enriched GO terms associated with the upregulated (orange) and downregulated (blue) genes. Each dot represents one gene. The color of the inner ring corresponds to the Z-score, wherein darker color indicates smaller Z-score. The height of the ring indicates the significance of the GO term (in terms of -log_10_ adj-p-value). (E) Chord plot showing the core genes associated with the significantly enriched GO terms among the downregulated genes. GO terms are shown on the right, and the genes are shown on the left. Logarithmic fold changes are denoted as gradient of blue rectangle representing each gene. This plot shows Sor regulates genes with interconnected functions. (F) Heatmap shows the expression of genes coding for the mitochondrial respiratory chain complexes after Sor-treatment.

### Sorafenib alters proteostasis of functionally interlinked protein clusters in muscle cells

As a next step, we employed an unbiased quantitative proteomic approach to map the protein complexes targeted in Sor-induced cachexia. Our quantitative mass spectrometry (MS) results identified that 61 proteins were upregulated, while 262 proteins showed reduced level (Fig. 5A), suggesting a net protein destabilizing environment in Sor-treated muscle cells. Similar to our transcriptomic data, the downregulated proteins in our proteomic screen predominantly belonged to GO terms of sarcomere organization, mitochondrial respiratory chain complex I, and sarcoplasmic reticulum, among others (Fig. 5B). STRING (Search Tool for the Retrieval of Interacting Genes/Proteins, version 12.0) network analysis identified a protein-protein interaction (PPI) network consisting of mitochondrial large (Mrpl1, Mrpl15, Mrpl48, etc.) and small (Mrps21, Mrps28, Mrps11, etc.) ribosomal subunits that are involved in mitochondrial protein translation (along with Gfm2 and Eif4a2) at a PPI enrichment p-value < 1.0e-16 (Fig. 5C). Another strongly enriched functionally interconnected cluster was formed by subunits of the mitochondrial respiratory chain complex I (Mt-Nd1, Mt-Nd5, Ndufv2, Ndufa12, etc.), which belongs to the downregulated category of proteins (Fig. 5D). We also observed significant downregulation of sarcomeric proteins, particularly troponin complex (Tnnt1, Tnni1, Tnnt3, etc.), sarcoplasmic reticulum (RyR1, Atp2a1) and T-tubule proteins (Cav3, Dysf), and proteins involved in actin binding (Synpo2, Nrap, Lmod2, etc.) and proteins involved in sarcomere organization (Myom3, Mybph, etc.) (Fig. 5E). A subset of the upregulated proteins showed overlap with our transcriptomic data (Fig. 4C), and were associated with ER-Golgi protein translocation and ERAD (ER-associated protein degradation) pathway (Hspa5, Sec61a, Herpud1, Ikbip, Cnih4, Cdipt, etc.) (Fig. 5A, Table S1-sheet 3). Overall, these observations indicate a high correlation between our transcriptomic and proteomic findings.

**Fig. 5:**
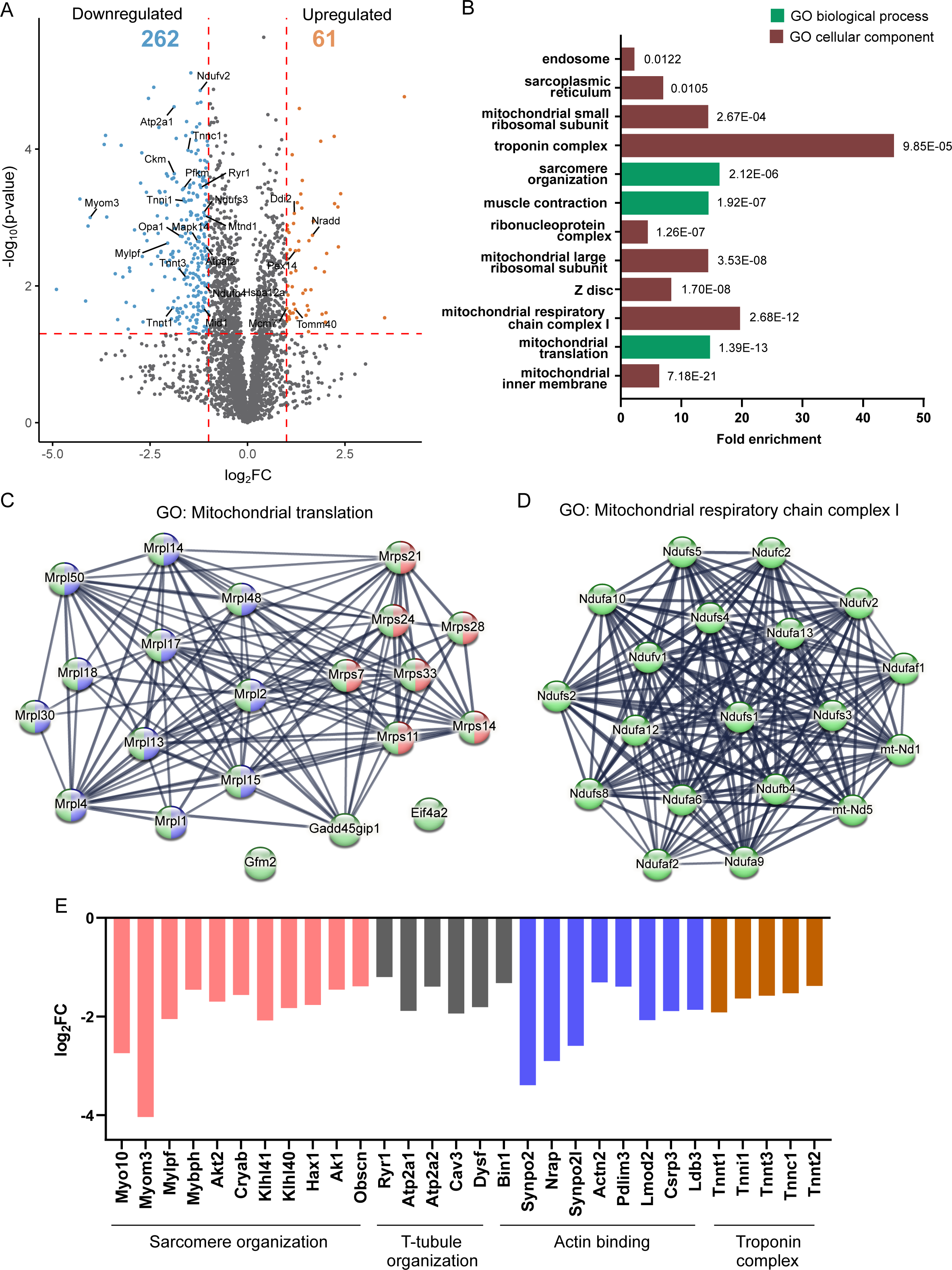
Quantitative proteomics identifies deregulated proteostasis-upon Sorafenib treatment-particularly of myofibrillar, SR and mitochondrial proteins. (A) Quantitative mass spectrometry (MS) results identified a total of 4634 proteins from independent MS screens across all samples. The proteins quantified in at least two out of three replicates (filtered with FDR < 0.01) were shortlisted, resulting in total 3452 proteins. The treatment groups (DMSO and Sor) were compared and proteins with log2-fold change ≥ ±1.0 and p-value ≤ 0.05, as indicated by the red dotted line, were labelled as significantly altered. This resulted in 323 proteins with differential expression (up- and down-regulated) upon Sor treatment. (B) Bar graph shows the GO terms enriched among the downregulated genes. Green indicates GO-biological process and brown indicates GO-cellular component. The head of each bar shows the adjusted p-value for that particular GO term, as estimated by DAVID analysis. (C) STRING network showing cluster of downregulated proteins associated with mitochondrial small (pink) and large (blue) ribosomal subunit, and mitochondrial translation (green). (D) Highly interconnected protein cluster determined by STRING analysis shows the downregulated protein subunits of the mitochondrial respiratory chain complex I. (E) Bar graph shows the expression of proteins associated with the following GO terms – orange= troponin complex, grey = T-tubule organization, blue = actin binding, pink = sarcomere organization.

To further validate our RNAseq and MS data, we performed additional independent experiments and probed for the transcript and protein expression of some of the key targets. In agreement with our RNAseq findings, we observed a significant reduction of ∼2 fold in *MyHC-IId and MyHC-IIa* transcript expression. In line with these results, we also detected reduced protein level of the motor protein MyHC-IId and MyHC-IIa (Fig. 6A). Noteworthy, the other two TKIs Nilo and Ima, which did not show any significant effect on sarcomere organization and muscle contractility, also did not alter gene expression of *MyHC-IId* or *MyHC-IIa* (Fig. S3A, B). Also, Sor treatment did not significantly change alpha-actinin transcript or protein expression (Fig. S3D and S1D).

**Fig. 6:**
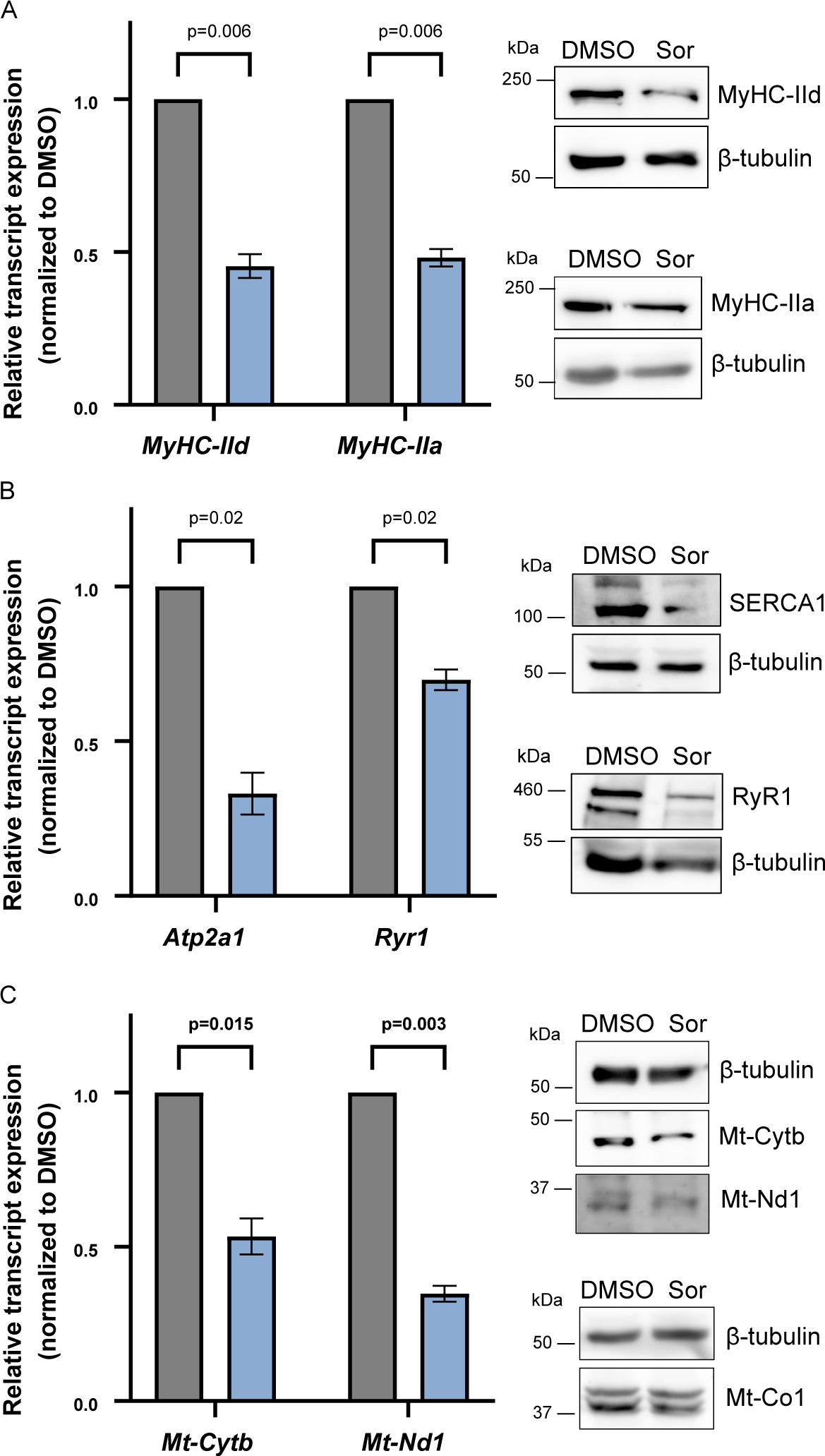
Sorafenib altered expression of major muscle-specific proteins. (A) Graph shows relative transcript expression of *MyHC-IId* and *MyHC-IIa* as measured by RTqPCR. Western blot shows protein expression of MyHC-IId and -IIa in DMSO- and Sor-treated myotubes. (B) Graph shows relative transcript expression of *Atp2a1* and *Ryr1*. Western blot shows protein expression of SERCA1 and RyR1 in DMSO- and Sor-treated myotubes. (C) Graph shows relative transcript expression of *Mt-Cytb* and *Mt-Nd1*. Western blot shows protein expression of Mt-Cytb, Mt-Nd1 and Mt-Co1 in DMSO- and Sor-treated myotubes. β-tubulin was used as the loading control for western blotting. The blots are the representative of at least three biological experiments. The transcript expression for each gene was normalized to *Gapdh* and its respective DMSO control. Data is represented as Mean ± SEM, from three biological replicates with technical triplicates in each experiments. Statistical significance was determined using unpaired t-test with Welch’s correction on GraphPad Prism 9.5.1. DMSO = grey, Sor = blue. Protein expression blots (B, D and F) are representatives of at least three biological experiments.

In our validation assays, we detected significant downregulation of SR genes *Ryr1* and *Atp2a1.* Corresponding to the transcriptional status, we observed a strong decrease in RyR1 and SERCA1 (gene - *Atp2a1)* protein level as well (Fig. 6B). To validate the genes associated with mitochondrial function, we focused on Mt-Nd1 (complex I) and Mt-Cytb (complex III) as our targets. Mt-Nd1 was one of the top downregulated complex I subunits in both RNASeq and MS datasets, while Mt-Cytb was specifically the only complex III subunit downregulated in our transcriptome data (Fig. 4F, 5D). Across the three experiments, we observed a significant reduction in transcript expression of Mt-Nd1 (∼2.9 fold) and Mt-Cytb (∼1.9 fold). The same was observed at the protein level (Fig. 6C). Noteworthy, another mitochondrial respiratory chain complex protein Mt-Co1 (complex IV) showed no apparent change in Sor-treated cells (Fig. 6C, right panel), highlighting target specificity by Sor. This was further substantiated by our RNAseq dataset, wherein the expression of OXPHOS complex II subunits (SDHA/B/C/D) remained unchanged upon Sor treatment. These observations indicate that Sor induces cachexia by specific deregulation of functionally interlinked protein clusters, primarily involved in sarcomere organization, muscle contraction, SR-mediated calcium handling and mitochondrial oxidative respiration. For the subset of tested candidate proteins, Sor affected the gene expression at transcriptional level, in addition to its effects on protein homeostasis, resulting in further reducing the protein amount of major players, including MyHC, SERCA1, Mt-Nd1 and Mt-Cytb, key for muscle cell metabolism and contractile function.

### Sorafenib regulates transcription by modulating epigenetic state of specific genes in muscle cells

The observed transcriptional deregulation may be a result of altered chromatin signalling, and serve as a prominent mechanism commencing Sor-induced cachexia. To test this idea, we optimized chromatin immunoprecipitation (ChIP) experiments and used *MyHC-IId* and *Atp2a1* as target genes. To this end, we designed promoter primer pairs to study possible chromatin-related changes triggered by Sor on these genes. Histone 3 lysine 4 trimethylation (H3K4me3) is one of the most prominent histone modifications that is typically associated with positive transcriptional activity ^49^. We measured H3K4me3 marks on these genes. DMSO- and Sor-treated myotubes were used for chromatin isolation. ChIP was performed with an antibody directed against H3K4me3. Additionally, a non-specific IgG antibody was used as an isotype control. The real-time qPCR experiments using ChIP-enriched DNA templates showed a strong enrichment of H3K4me3 over the IgG control (∼33.6 fold) on the *MyHC-IId* promoter, highlighting the specificity of our ChIP assays. The abundance of H3K4me3 was significantly reduced by ∼3 fold upon Sor treatment (Fig. 7A). Similar results were observed for *Atp2a1*, whereby Sor treatment significantly reduced H3K4me3 by ∼2.7 fold on the gene promoter (Fig. 7A). These outcomes suggested that Sor altered muscle-specific gene expression by reducing epigenetic marks associated with active transcription.

**Fig. 7:**
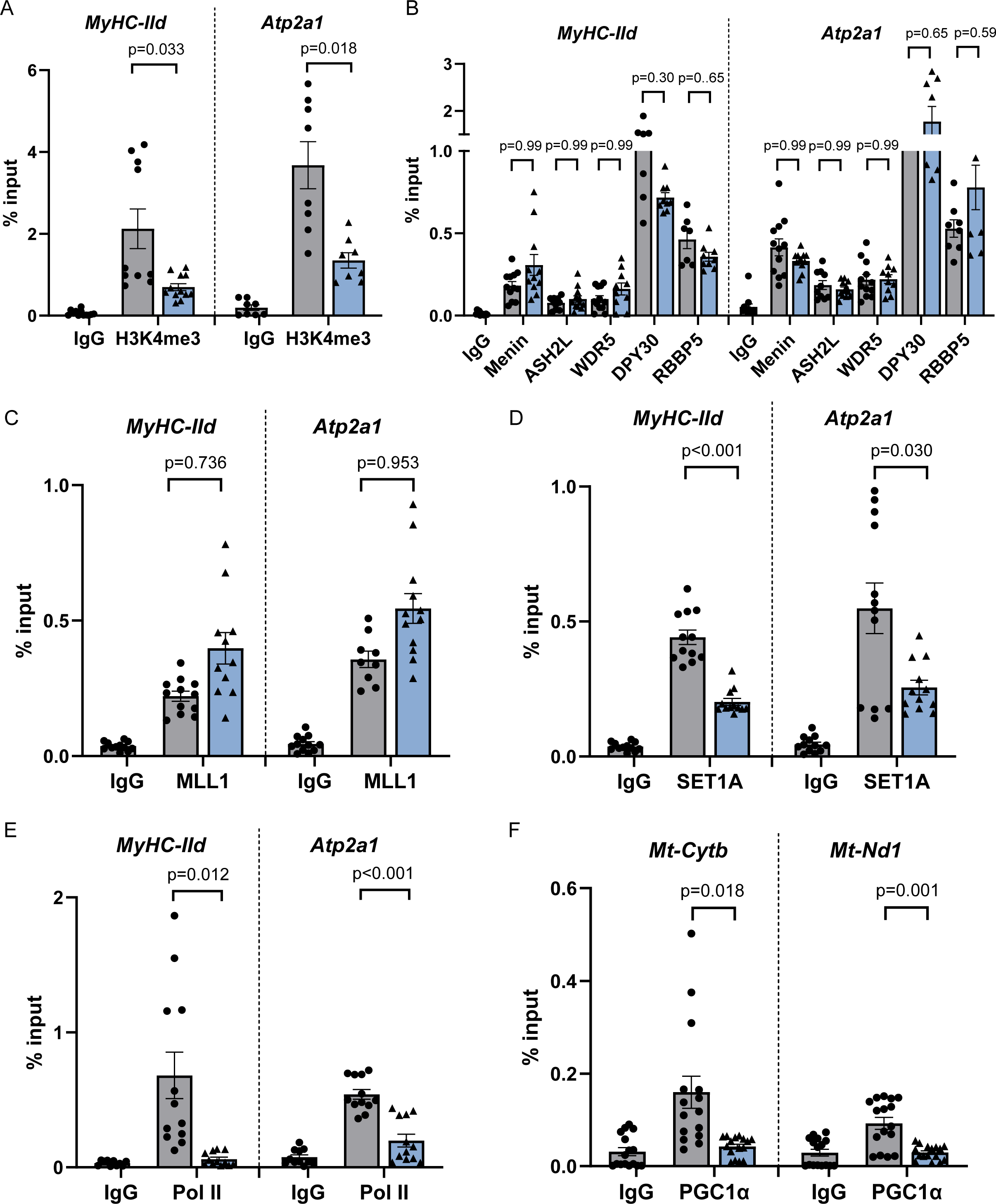
Sorafenib deregulates chromatin signalling of muscle-specific genes. ChIP experiments show chromatin enrichment of (A) H3K4me3, (B) SET1/MLL complex proteins – Menin, ASH2L, WDR5, DPY30 and RBBP5, (C) catalytic core MLL1 and (D) SET1A on *MyHC-IId* and *Atp2a1* gene promoters, (E) Similar to Fig. 7A, except anti-RNA Pol II use used for ChIP assays. The ChIP data represent the Mean ± SEM from three biological replicates with technical quadruplicates. Except DPY30 and RBBP5 ChIPs, where N= 2 and n =8 (N= biological replicates, n= technical replicates) (F) Graph shows chromatin association of PGC1α on the promoter of mitochondrial genes *Mt-Cytb* and *Mt-Nd1*. N= 3 and n =12 (N= biological replicates, n= technical replicates). IgG was used as the antibody isotype control. Statistical significance was determined using unpaired t-test with Welch’s correction on GraphPad Prism 9.5.1. DMSO = grey, Sor = blue.

### Sorafenib specifically downregulates chromatin association of SET1A histone methyltransferase complex

SET1/MLL complex is the major epigenetic modifier that adds methyl groups on H3K4 (H3K4me3) of its target genes. Presence of this complex on its target gene can influence the gene activity ^31^. Since Sor specifically reduced H3K4me3 on distinct muscle-specific genes, we asked if Sor might affect SET1/MLL complex chromatin association. To test this possibility, we first checked if the SET1/MLL complex directly regulates the candidate genes *MyHC-IId* and *Atp2a1.* ChIP assays revealed a strong enrichment of WRAD subunits WDR5, RbBP5, ASH2L and DPY30 on these genes as compared to IgG (Fig. 7B). The catalytic core MLL1 itself and the MLL1/2-specific subunit Menin was also enriched, suggesting our candidate genes are direct targets of the SET1/MLL complex (Fig. 7B, C). We found no significant change in chromatin occupancy of the WRAD subunits and the catalytic core MLL1 on the target gene promoters (Fig. 7B, C). Despite the unchanged chromatin occupancy of MLL1 catalytic core and the WRAD complex, significant reduction in H3K4me3 marks was observed on these genes (Fig. 7A). We reasoned if Sor modulates other catalytic core proteins and therefore, examined another catalytic core protein SET1A. ChIP using anti-SET1A antibody showed a strong chromatin enrichment of SET1A over IgG on *MyHC-IId* (∼12 fold) and *Atp2a1*(∼12.2 fold) promoters in DMSO-treated cells (Fig. 7D), further confirming these genes as direct targets of SET1A. Interestingly, compared to control, Sor-treated cells revealed significantly reduced chromatin association of SET1A on *MyHC-IId* (∼2 fold) and *Atp2a1* (∼2 fold) gene promoter (Fig. 7D). These findings strongly indicated that Sor specifically alters the epigenetic marks on *MyHC-IId* and *Atp2a1* by influencing chromatin residency of SET1A, thereby resulting in decreased transcriptional output. Finally, to investigate if the lowered H3K4me3 indeed decreased the transcriptionally active state, we probed the chromatin association of RNA polymerase II (Pol II) on these genes. Our ChIP assays revealed a significantly reduced association of transcriptionally active Pol II on *MyHC-IId* (∼11 fold) and *Atp2a1* (∼2.8 fold) gene (Fig. 7E), suggesting that Sor reduced transcriptional competency of these genes.

Next, we focused on mitochondrial-encoded candidate genes *Mt-Nd1* and *Mt-Cytb,* which were altered by Sor treatment. Peroxisome proliferator-activated receptor-gamma coactivator (PGC)-1α is a transcriptional co-activator that regulates the expression of genes encoded by the mitochondrial DNA (mtDNA), namely the OXPHOS proteins ^50^. We investigated if Sor influences PGC1α association via ChIP experiments. As compared to IgG, we obtained strong enrichment of PGC1α on the target gene promoters (∼3 to 5 fold). This enrichment was significantly reduced on *Mt-Cytb* (∼3.8 fold) and *Mt-Nd1* (∼3.14 fold) upon treatment with Sor (Fig. 7F). These observations suggest that Sor alters the PGC1α-mediated transcription of mitochondrial respiratory chain complex proteins. Taken together, Sor-targeted chromatin events of distinct nuclear- and mitochondrial genome-encoded genes, resulting in transcriptional deregulation that ultimately induced cachexia.

### Sorafenib induces cachectic phenotype in satellite cell-derived primary muscle cells

To further strengthen our findings, we validated some key observations in murine satellite cell (MuSc)-derived primary muscle cells. To this end, we harvested MuSCs from mice, differentiated into myotubes and treated with Sor. We observed significant downregulation of myosin heavy chain *MyHC-IId* and *MyHC-IIa* (∼2.8 fold) at the transcript level upon Sor treatment (Fig. 8A). The expression of *alpha-actinin* transcript remained largely unaltered. Additionally, transcript expression of SR Ca^2+^-handling proteins (*Atp2a1* and *Ryr1)* and mitochondrial respiratory chain complex subunits *(Mt-Nd1* and *Mt-Cytb)* was significantly reduced (Fig. 8B, C). Furthermore, consistent with our observations from the cell culture model, confocal imaging of Sor-treated primary muscle cells revealed severe disruption of the sarcomere assembly, in contrast to the control cells that showed a well-resolved, striated sarcomeric pattern (Fig. 8D and Fig. S4). Collectively, these results substantiated our findings that Sor causes cachexia by deregulating muscle-specific gene expression program.

**Fig. 8:**
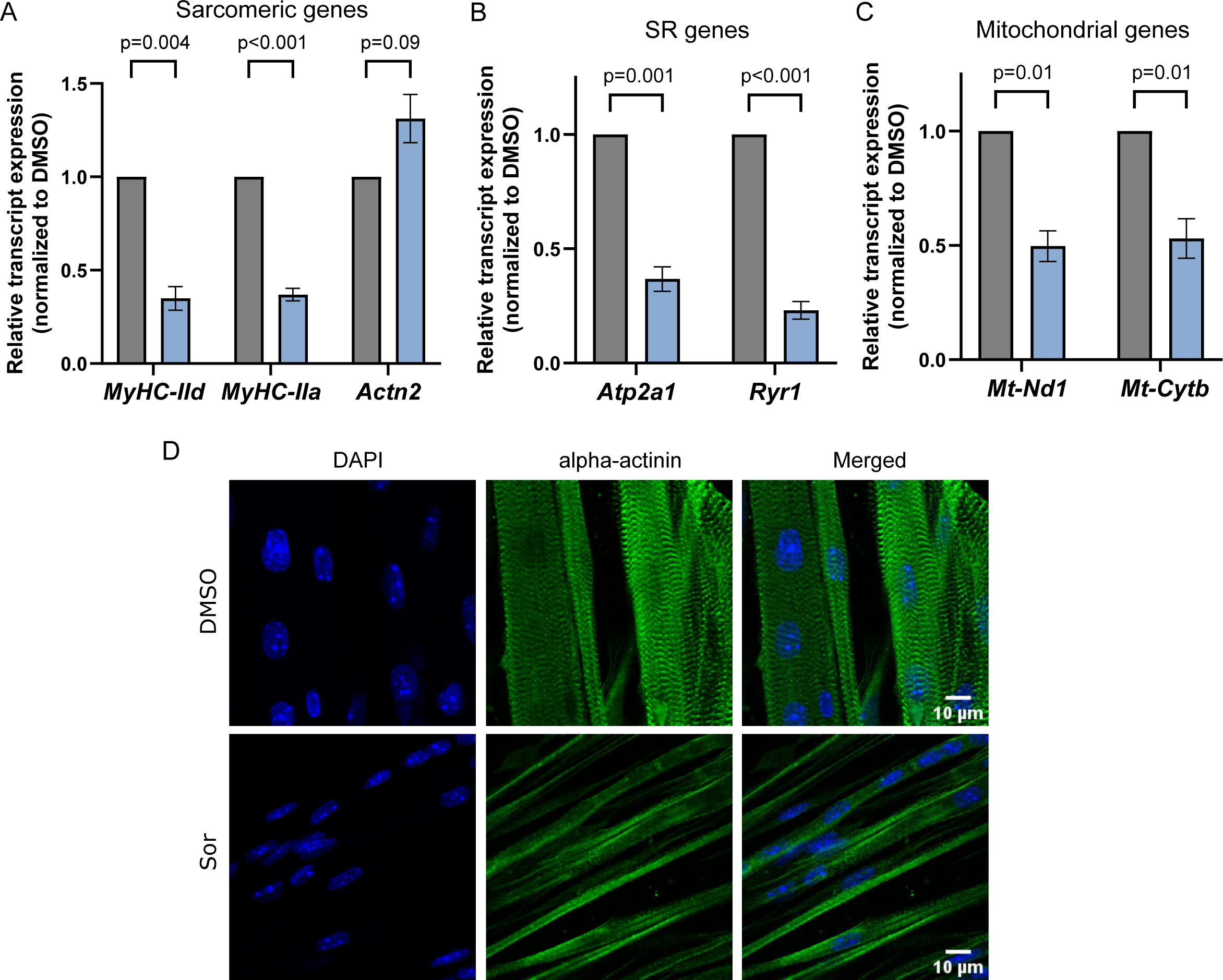
Effects of Sor in satellite cell-derived muscle cells. Graph shows relative transcript expression of (A) sarcomeric genes – *MyHC-IId*, *MyHC-IIa* and *Actn2*, (B) SR Ca^2+^-handling genes – *Atp2a1* and *Ryr1*, and (C) mitochondrial OXPHOS genes – *Mt-Nd1* and *Mt-Cytb*, in DMSO- and Sor-treated primary muscle cells. The transcript expression for each gene was normalized to *Gapdh* and its respective DMSO control. All graphs show Mean ± SEM, where N = 4, n = 12. (N= number of mice, n= technical replicates). Statistical significance was determined using unpaired t-test with Welch’s correction on GraphPad Prism 9.5.1. DMSO = grey, Sor = blue. (D) Panel shows representative confocal microscopy images of satellite cell-derived muscle cells immunostained against alpha-actinin (green) and nucleus (DAPI – blue). Images were captured using the 40X objective. Scale bar = 10 µm. See also Fig. S4. The images are representative of primary muscle cells derived from a total of seven different mice.

### Sorafenib hinders myogenic differentiation and myoblast proliferation

Myogenic differentiation process is critical for generating mature functional muscle cells as well as repair of existing muscle fibres ^51–53^. As shown in earlier sections, we observed muscle wasting and loss of muscle function due to Sor-induced alterations in distinct interconnected pathways; we further probed if Sor can affect myogenic differentiation process. To this end, we calculated the myogenic index of myoblasts treated with Sor vs DMSO. Immunostaining against sarcomeric protein alpha-actinin revealed that fewer myotubes developed from Sor-treated myoblasts (Fig S5A). Correspondingly, the myogenic index of Sor-treated myoblasts (6%) was significantly lower as compared to control myoblasts (68%) (Fig. S5A). When probed for the expression of key myogenic regulatory factors (MRFs) in these cells, we observed significantly low protein expression of MEF2C and MyoD, signifying an impaired myogenic differentiation ability of myoblasts upon treatment with Sor (Fig. S5B). In line with the above observations, cells also showed strong reduction in expression of myofibrillar proteins MyHC-IId and IIa (Fig S5C). In addition, Sor also severely compromised myoblast proliferation as the percentage of Ki67-one of the marker for proliferation-positive cells was reduced by approximately 50% upon Sor treatment, as compared to control cells (Fig S5D). Taken together, Sor induce cachexia by hampering myogenic differentiation as well as perturbing the intracellular organization, cell metabolism, and consequently the function of mature muscle cells.

## Discussion

In this study, we identified a system-wide reprogramming of interconnected physiological pathways in Sor-induced cachexia. Sarcomere organization, calcium homeostasis and mitochondrial dynamics in mature muscle cells were the primarily altered cellular processes. Although the proteomic screen revealed that Sor directly influences stability of a large number of proteins (for example, mitochondrial ETC complex IV subunits-Coq5, Cox20), it is likely that the observed Sor-mediated transcriptional deregulation of distinct muscle-specific genes is the upstream event. Our in-depth investigations uncovered that these deregulated nuclear genes are targets of SET1/MLL histone methyltransferase complex. Sor reduced the chromatin association of SET1A, one of the catalytic core proteins of the SET1/MLL complex. Consequently, the classical epigenetic mark of productive transcription i.e., H3K4me3 was diminished from these genes. Thus, it is conceivable that the modulation of chromatin-related events by Sor is the nucleation event in Sor-induced cachexia. Although few studies ^54,55^ have indicated a potential link between Sor and muscle wasting, a comprehensive understanding of the implications of Sor in muscle cell organization and function remained elusive. Our study delineated distinct molecular pathways derailed due to Sor-treatment, leading to cachectic phenotype. Importantly, we also found a reduced differentiation potential of muscle progenitor cells upon Sor treatment, thus reinforcing the damaging influence of Sor on muscle functions.

The observed Sor-induced loss of muscle cell contractility could be attributed to the multipronged effects of Sor, which was revealed by our system-wide analysis as well as target-based investigation. Sor not only affected sarcomere organization and cell contractility, but it also impaired SR-mediated calcium handling and reoriented mitochondrial metabolism leading to insufficient energy production. The diminished transcriptional activity of specific sarcomeric genes, particularly the myosin heavy chain isoforms *MyHC-IId* and *MyHC-IIa* directly affected the contractile potential in Sor-treated cells. The hexameric myosin protein is composed of two heavy chains and four light chains i.e., regulatory (RLC) and essential light chain (ELC). As a major force generator, the individual components of the myosin complex are key to its proper functioning ^56^. Besides reduction in *MyHC,* our transcriptomic dataset showed reduced expression of the ELCs (gene-*Myl1*) following Sor treatment (Fig. 4C, E). These findings suggests that Sor targeted a specific group of functionally interconnected sarcomeric genes. Besides, Sor reduced the transcriptional output of various SR genes and thereby proteins involved in calcium handling (such as *Ryr1* and *Atp2a1*), resulting in absence of calcium transients. A recent study reported a moderate effect on the calcium transients in Sor-treated mouse ventricular cardiomyocytes, where Sor effect was relayed through reduced phosphorylation of phospholamban protein ^57^. Here, we observed a complete abrogation of calcium transients upon Sor treatment in skeletal muscle cells. We attributed Sor-induced defective calcium handling to reduced expression of major Ca^2+^-binding proteins as validated with our transcriptomic and proteomic measurements (Fig. 4E, 5E). Our findings unravelled a distinct mechanism of Sor-induced cachexia in skeletal muscle cells. It seems probable that since the skeletal muscle and cardiomyocytes share similar structural and functional features, the nature and extent of effects might be identical. However, our results indicate that the mechanism by which Sor targets calcium handling in cardiomyocytes may be somewhat different from in skeletal myocytes. Since TKI can induce cardiotoxicity ^58^, further comparative studies may be required to address this question.

In addition to deregulated SR function, we also observed impairment of mitochondrial function in Sor-treated cells, characterized by changes in mitochondrial membrane potential, that ultimately hampered ATP production in Sor-treated muscle cells. Sor-treated cells also displayed significantly lower oxidative respiration via the ETC and a shift towards the energy-inefficient glycolytic mode of respiration (Fig. 3D-G). A recent report ^59^ showed impaired mitochondrial oxidative respiration in Sor-treated human induced pluripotent stem cell-derived cardiomyocytes (hiPSC-CMs). However, the relevance of this observation in skeletal muscle and the underlying molecular mechanism of this observation was not clearly known. Our muscle cell system-wide study provided a detailed molecular insight of Sor-induced alteration of cellular respiration. The ChIP assays identified the origin of Sor-induced defective mitochondrial function to be decreased chromatin association of PGC1α, the master transcriptional regulator of mitochondrial gene expression and mitochondrial biogenesis. Chromatin association of PGC1α was reduced for OXPHOS pathway genes *Mt-Nd1* and *Mt-Cytb*, resulting in decreased expression of OXPHOS complex I and III subunits, respectively. This regulation led to lower gene and protein expression of Mt-Nd1 and Mt-Cytb. The same was in agreement with our transcriptomic and proteomic analyses (Fig. 4F, 5D). Apart from transcriptional deregulation, Sor directly influenced mitochondrial protein homeostasis by reducing the protein expression and/or stability of mitochondrial small (Mrps) and large (Mrpl) ribosomal subunits, as detected in our proteomic screen (Fig. 5C). This suggests an insufficiency in the assembly and number of mitochondrial ribosomes, which when coupled to a reduction in expression of other mRNA translation factors (Gfm2 and Eif4a2), ultimately resulted in impaired mitochondrial protein synthesis. Interestingly, a significant downregulation of Creatine kinase (Ckm, an enzyme involved in energy production) in our RNAseq experiment additionally provided evidence of reduced energy production capacity by mitochondria upon Sor treatment (Table S1). Taken together, an unprecedented Sor-mediated defect on molecular pathways governing mitochondrial dynamics and cellular respiration was shown.

Interestingly, Sor treatment significantly enhanced the expression of proteins associated with protein folding responses of the endoplasmic reticulum (ER) and Golgi complex such as PERK, ATF6, IRE1, BiP and ATF4 (Table S1-sheet3). We also observed a strong upregulation of molecular chaperones such as Hsp70 in both transcriptomic and proteomic datasets. Our observation corroborates previous findings where these proteins, particularly Hsp70, were reported to be upregulated and activated in atrophic environment associated with cachexia ^60^. Noteworthy, many important regulators of sarcomere organization, SR and mitochondria remained unchanged after Sor treatment (Table S1), highlighting a specific effect of Sor in muscle cells.

A remarkable finding in this study is the effect of Sor in alteration of H3K4me3 epigenetic marks on muscle-specific RNA polymerase II (Pol II)-regulated nuclear genes, particularly *MyHC-IId* and *Atp2a1*. Our results suggest that diminished H3K4me3 is most likely the main reason of lower transcriptional output of Sor-responsive gene promoters. In line with this observation, we found the chromatin association of the “writer” of H3K4me3 (i.e., the SET/MLL complex) was also affected in Sor treatment. Noteworthy, the chromatin association of only SET1/A-but not other MLL complexes and its subunits that we have tested here-was affected. Thus, Sor affected specific epigenetic program of distinct genes towards inducing cachexia. Previously, various other non-TKI class drugs such as daunorubicin (Daun) and VP16 etoposide were shown to have additional mechanisms of action than inhibiting topoisomerases. In cancer cells, these drugs can target chromatin domains by sensing specific epigenetic marks and induce DNA damage of these specific chromatin domains ^61^. Currently, we do not know if these mechanisms are also common in Sor-induced cachexia. However, these observations underscore the unknown molecular pathways regulated by various kinds of drug that possibly work in a cell-type-dependent manner. In our current study, we showed an unexpected link between the Sor-regulated function of the SET1/MLL complex, which led to altered H3K4me3 on SET1/MLL complex target genes in muscle cells. To the best of our knowledge, molecular insights regarding the role of Sor with TrxG family of epigenetic regulators (SET1/MLL) and associated epigenetic processes, particularly the H3K4me3, was not previously determined in muscle cells. Our study filled this gap and provided in-depth information on the cause of the cellular and functional phenotype.

Another striking finding of this study is the differential effect of the three different TKIs tested here. While Sor induced cachexia, the other two drugs Nilo and Ima showed no discernible effects. Unlike Sor, both Nilo and Ima did not alter muscle cell contractility, and neither did they perturb sarcomere organization. In addition, Nilo and Ima also did not deregulate the expression of genes that were downregulated by Sor (Fig. S3). This indicates that Sor-mediated mechanism seems to be not conserved among other tested TKIs such as Nilo and Ima. Thus, our work provides a framework for careful investigation of TKIs that might lead to the development of improved non-toxic cancer treatment regimens. This might blunt chemotherapy-induced cachexia and would enhance life quality and expectancy. Currently, we do not know whether Nilo and Ima might be involved in global transcriptional regulation and proteostasis in muscle cells. Future investigations are necessary to test these aspects.

In conclusion, this study highlighted the differential effects of the selected TKI family of drugs on muscle function. Our findings provide a conceptual understanding of Sor-induced cachexia by revealing detailed mechanistic insights on Sor-modulated epigenetic processes and the resulting transcriptional deregulation of distinct muscle-specific genes. These genes govern diverse but functionally interconnected pathways in muscle cells. The correct choice of drugs with minimal side effects or potentially damaging effects that can be countered is possible only with the knowledge of underlying affected pathways. Thus, the implications of these findings are directly relevant for developing balanced combination therapies for affected individuals to improve the treatment.

## METHODS

### List of antibodies

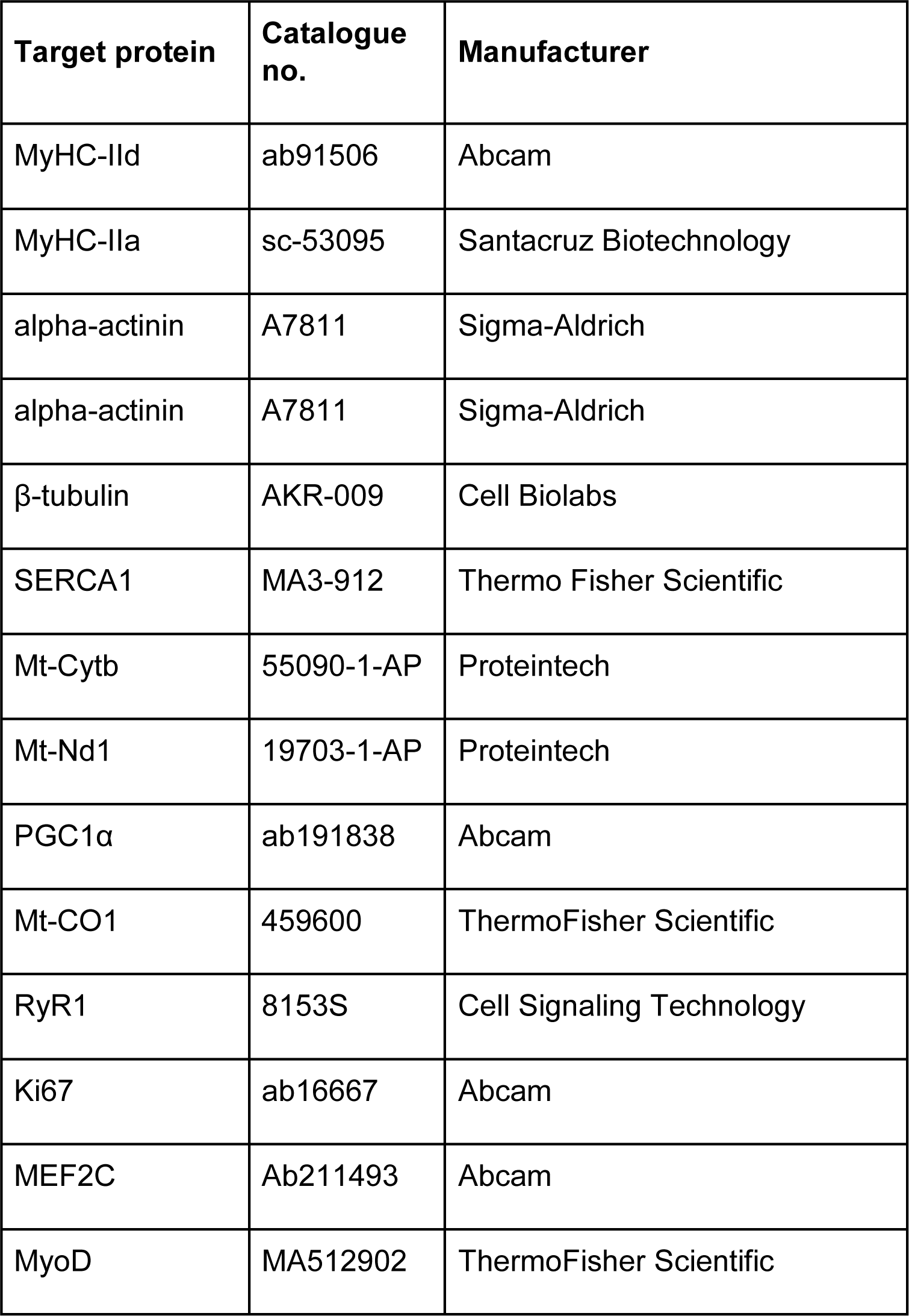

### List of oligonucleotides

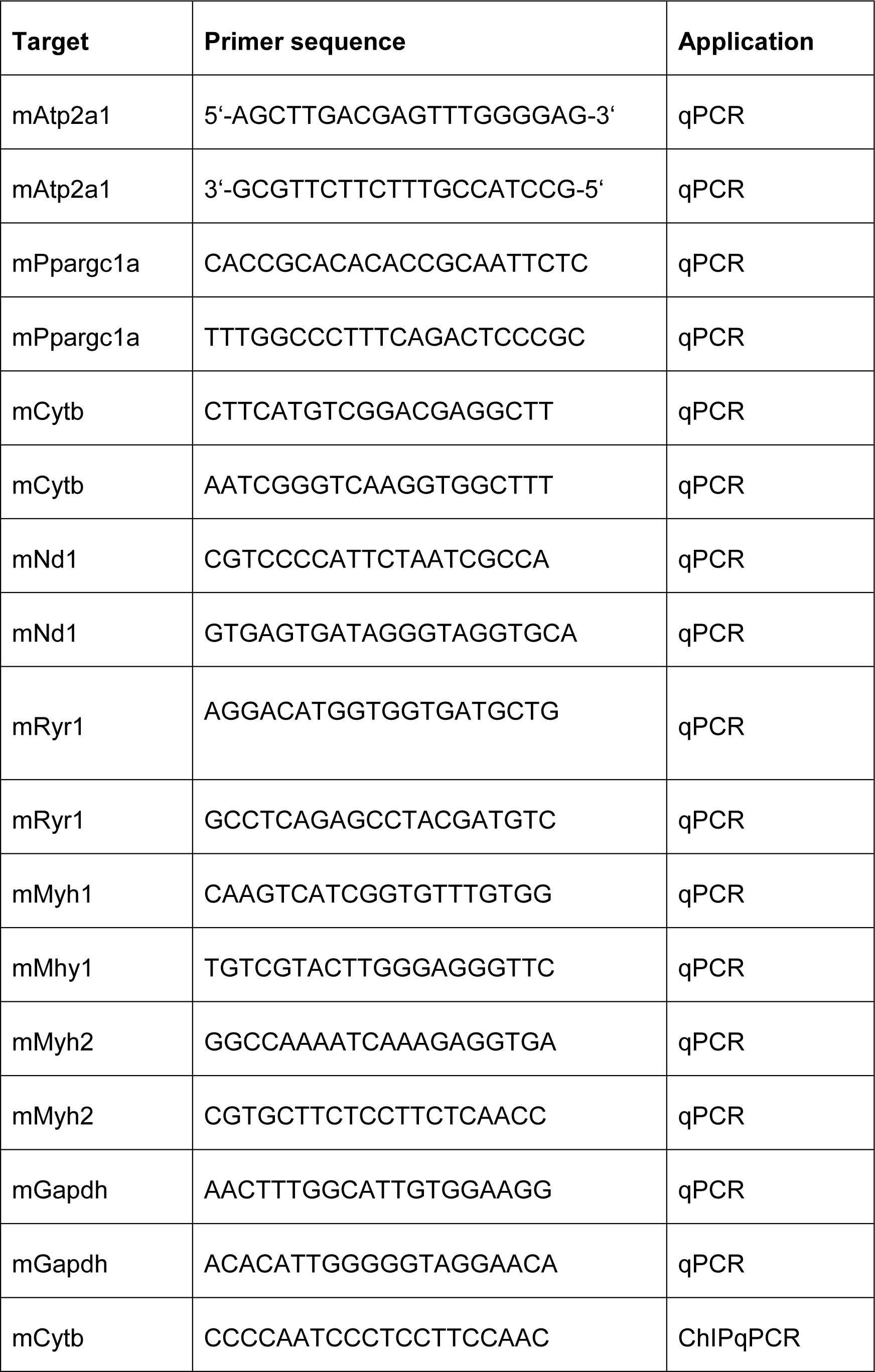
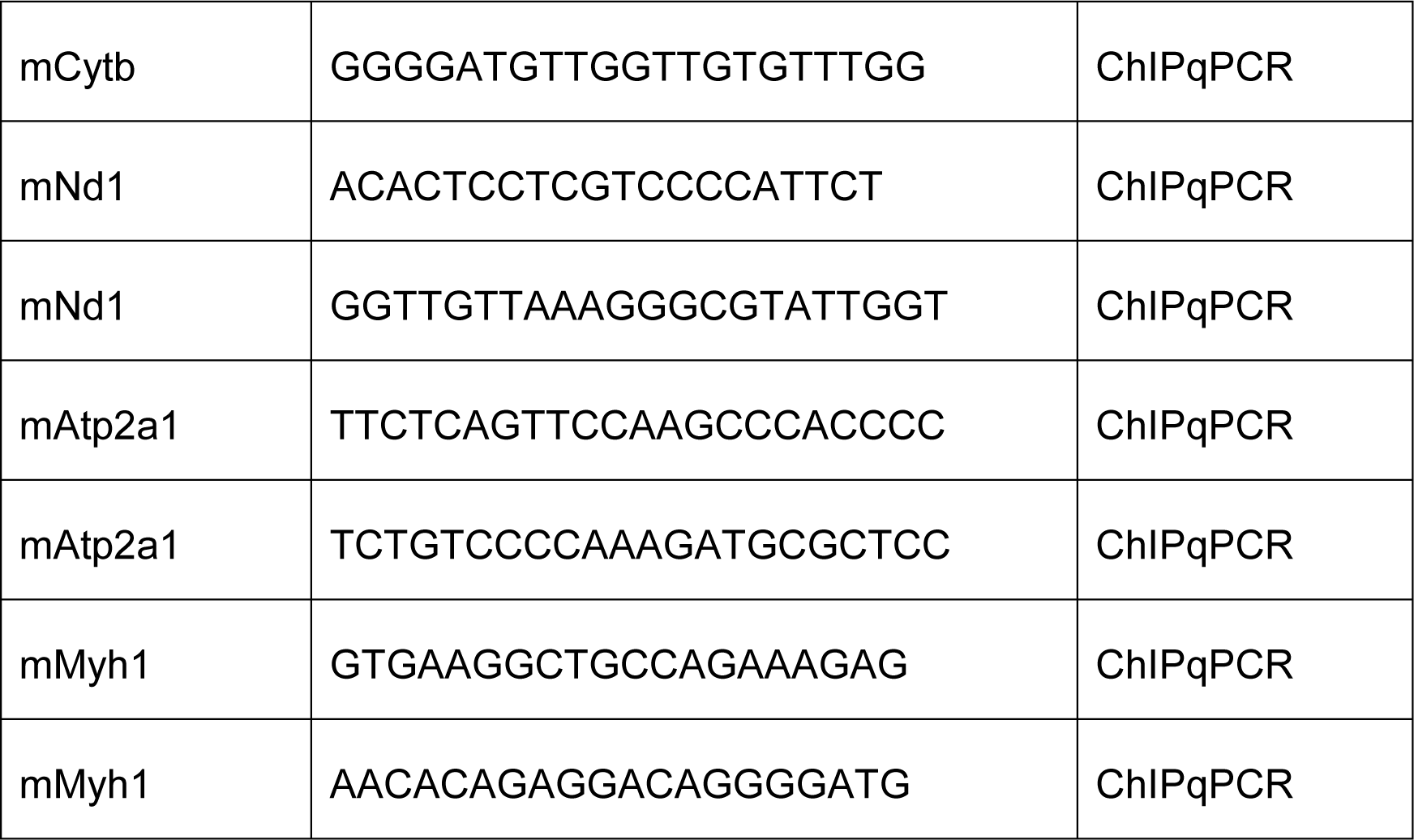

### Cell culture and drug treatment

C2C12 mouse myoblast cells were cultured in High Glucose DMEM (#DMEM-HA, Capricorn Scientific GmBH, Germany) supplemented with 15 % heat inactivated fetal bovine serum (#10100-147, Thermo Scientific, US) and 1 % Penicillin/Streptomycin (#PS-N, Capricorn Scientific GmBH, Germany). Once the cells reached ∼90% confluency, differentiation was induced using High Glucose DMEM supplemented with 2 % heat inactivated horse serum (#26050-088, Thermo Scientific, US) and 1% Penicillin/Streptomycin, with fresh medium being replaced every 24 h to generate mature myotubes. On day 4 of differentiation, the myotubes were treated with tyrosine kinase inhibitors (TKIs) – Sorafenib (SML2653-5MG, Sigma-Aldrich, Germany), imatinib mesylate (S1026, Selleck Chemicals GmbH, Germany) and nilotinib (12209, Cell Signaling Technology, Germany) at an assay-dependent concentration for 24 h. Corresponding volume of DMSO was used as vehicle control. For treatment of myoblasts, the cells were seeded at an assay-dependent cell density. After overnight incubation, the cells were treated with 20 µM Sor and DMSO and incubated for 24 h. After removal of the drug, the cells were allowed to proliferate for additional 24 h, then either differentiated into myotubes or harvested, based on the experimental set-up.

### Satellite stem cell isolation, differentiation and treatment

Male and female wild type C57BL/6J mice were bred in a certified facility at San Raffaele Hospital, Milan, Italy (authorization n. N. 11/2022-PR) and sacrificed 1-2 months after birth. The experimental procedures involving mice were conducted in accordance with animal welfare laws and guidelines approved by the Italian Ministry of Health and the Institutional Animal Care and Use Committee. Hind-limb muscles were collected from sacrificed mice and stored at 4-8°C for maximum 1 h before dissociation. Muscles were minced to 1 mm pieces and digested for 45 min at 37°C in 1X PBS (#ECB4004L, Euroclone, Italy) supplemented with 2.4 U/ml dispase II (#4942078001, Roche, Switzerland), 2 mg/ml collagenase A (#1013586001, Roche, Switzerland), 0.2 mM CaCl_2_ (#C5670, Sigma-Aldrich, Germany), 4 mM MgCl_2_ (#M8266, Sigma-Aldrich, Germany) and 0.1 mg/ml DNase I (#1014159001, Roche, Switzerland). The samples were resuspended in ice-cold Hanks balanced salt solution (HBSS) (#14025-050, Gibco, US) implemented with 0.2 % BSA and spun down at 340 rcf. The cell pellet was resuspended in HBSS containing 0.2 % BSA, 1 % DNase I, 1 % PenStrep (#ECB3001, Euroclone, Italy) and dispersed by passing through a 70 µm and then 40 µm cell strainer, followed by an additional wash of the filter. Finally, the cells were spun down, resuspended in 300 µl ice-cold HBBS and stained with the following antibodies for 45 min:-PB-CD45 1:50 (#48-0451-82, eBioscience, US), PB-CD31 1:50 (#48-0311-82, eBioscience, US), PB-Ter119 1:50 (#48-5921-82, eBioscience, US), FITC-Sca1 1:50 (#11-5981-82, eBioscience, US), and APC-α7integrin 1:100 (#67-001-05, AbLab, Canada). We sorted MuSCs by using BD FACS ARIA III SORP as PB-CD45^−^/PB-CD31^−^/PB-Ter119^−^/FITC-Sca1^−^/APC-α7integrin^+^. The MuSCs were counted with Burker counting chamber and plated on 12-or 24-well dishes or on chamber slides (#80826-G500, Twin Helix) coated with 1 mg/ml matrigel (#354234, Sigma-Aldrich, US) at a seeding density of 5000 cells/cm^2^ and allowed to grow in DMEM (#10569-010, Gibco, US) supplemented with 20 % FBS (#35-015-CV, Corning, US), 10 % HS (#26050-088, Gibco, US), 1 % Pen/Strep, 1% chicken embryo extract (#CE650-DL, Seralab, UK) and 2.5 ng/ml bFGF (#13256029, Gibco, US) for 5 days, changing medium every 2 days. After the cells reached ∼90 % confluency, they were differentiated into myotubes using DMEM supplemented with 5 % HS and 1 % Pen/Strep for 4 days. Differentiated myotubes were treated with 20 µM Sorafenib for 24 h. Corresponding volume of DMSO was used as vehicle control.

### Myotube contractility assay

C2C12 myotubes differentiated for four days on 10 mm laminin-coated coverslips were treated with 0.1 µM nilotinib (Nilo), 20 µM imatinib (Ima) and 20 µM Sorafenib (Sor) for 24 h. On day 5, the coverslips were placed in a custom-designed glass chamber containing ∼300 µl pre-warmed medium. The chamber was connected to the mTCII micro-temperature controller (IonOptix, US) to maintain the temperature at 37°C. The apparatus was further connected to the MyoPacer field stimulator (IonOptix, US) that provides electrical stimulation at 40 V and 1 Hz. Using the 10X objective of the Olympus IX51 bright field microscope coupled to the Olympus XC30 camera (Olympus LifeScience, Japan), multiple videos were recorded at 10 fps (frame per second) to capture the myotube contractility. The number of contracting myotubes was manually counted and kymographs were generated using the ImageJ software.

### Intracellular calcium transient measurements

Sor-treated myotubes on 18 mm laminin-coated coverslips were loaded with 5 µM Ca^2+^ indicator FURA-2 AM (#F1221, Thermo Scientific, US) along with 2.5 mM Probenecid (#P36400, Thermo Scientific, US) and 0.025 % (w/v) Pluronic F-127 (#P6866, Thermo Scientific, US) in serum-free medium for 1 h at 37°C. The cells were washed twice with medium for 15 min each before recording. The coverslip was mounted on a custom-designed perfusion chamber connected to the MyoCAM setup (IonOptix, US) and perfused with buffer containing 20 mM HEPES, 1.2 mM NaH_2_PO_4_, 0.66 mM MgSO_4_, 117 mM NaCl, 5.7 mM KCl, 5 mM Na-Pyruvate, 1.25 mM CaCl_2_, 10 mM Creatin and 10 mM Glucose, set to pH 7.4. The cells were paced at 30 V and 1 Hz at 37°C maintained by the NBD TC2 Bip temperature controller (Cell MicroControls, US). The 40X objective of the Olympus IX71 microscope (Olympus LifeScience, Japan) was used to collect single cell data. Using a dual-excitation photomultiplier system interface (IonOptix, US), the emitted fluorescence was recorded at 510 nm after excitation at 340 and 380 nm to detect bound and free Ca^2+^ respectively. Background fluorescence from non-FURA-treated myotubes was used for normalization. The IonWizard software (IonOptix, US) was used to analyze the data.

### SDS-PAGE and Western Blotting

Whole cell lysates (WCL) were prepared using 4X ROTILoad buffer (#K929.1, Carl Roth, Germany) and DPBS in 1:4 ratio, followed by heating at 92°C for 10 min. Samples were loaded alongside the Precision Plus protein ladder (#1610394, Bio-Rad Laboratories Inc., US) on 10 % SDS-polyacrylamide gel, and run at 150 V for 70-80 min. The resolved proteins were transferred on to a 0.22 µ nitrocellulose membrane (#10600004, Cytiva, US) for 60 min at 10 V using the Trans-blot semi-dry transfer apparatus (Bio-Rad Laboratories Inc., US). The blots were blocked with 4 % skimmed milk (#2325, SantaCruz Biotechnology Inc., US) for at least 60 min at room temperature, and probed overnight with the respective primary antibodies, followed by one hour incubation with the secondary antibody. Post chemiluminescent development (#34075, Thermo Scientific, US), the blots were imaged using ImageQuant LAS4000 (GE Healthcare Technologies Inc., US).

### Immunofluorescence and confocal microscopy

Myotubes cultured on laminin-coated coverslips were fixed using 4 % PFA for 20 min, followed by permeabilization using 0.2 % (v/v) Triton X-100 for 10 min. Standard immunostaining protocol was followed and the coverslips were mounted on glass slides using fluoroshield (#F6182, Merck, Germany). Images were captured using the 10X and 60X oil immersion objective of the Olympus FluoView 1000 confocal microscope (Olympus LifeScience, Japan) at the Research Core Unit for Laser Microscopy, Hannover Medical School, Germany.

### Reverse transcription-quantitative polymerase chain reaction (RT-qPCR)

Total RNA was isolated using the High Pure RNA Isolation kit (#11828665001, Roche, Switzerland), followed by cDNA synthesis with the Maxima First Strand cDNA synthesis kit ((#A25741, Thermo Scientific, US) using 0.7 µg RNA as template. Quantitative PCR was performed at the QuantStudio 6 Flex real-time PCR system (#4485691, Thermo Scientific, US) to determine the transcript expression of target genes using the respective primer pair and SYBR green master mix (#A25741, Thermo Scientific, US). *Gapdh* was used as the housekeeping gene for normalization. Relative transcript expression was calculated using the ΔΔC_T_ method.

### Seahorse XF Cell Mito Stress Test

Myotubes were cultured in 96-well Seahorse XF cell culture microplates (#103015-100, Seahorse Bioscience, Agilent Technologies, US). Before running the assay, the cells were incubated with Seahorse XF DMEM assay medium (#103680-100, Seahorse Bioscience, Agilent Technologies, US) for 1 h at 37°C in absence of CO_2_. Using the same medium, 10X solutions of Oligomycin (OM) (port A), FCCP (port B) and Rotenone/Antimycin A (R/AA) (port C) were prepared and loaded on the cartridge plate. The XFe96 analyzer (Seahorse Bioscience, Agilent Technologies, US) sequentially injects the compounds from the cartridge plate into the cell culture microplate and records the changes in medium O_2_ and pH. The OCR and ECAR values were normalized with the total protein content (measured using Bradford assay) of each sample using the Seahorse Wave 2.6.3 software. Based on the OCR values, several assay parameters were calculated as follows-(i) Basal respiration = [OCR value of last measurement before OM addition] – [OCR value of last measurement after R/AA addition], (ii) Coupling efficiency = [(OCR value of last measurement before OM addition) – (Minimum OCR value after OM addition)] / [basal respiration] * 100, (iii) Spare respiratory capacity = [(Minimum OCR value after FCCP addition) – (OCR value of last measurement after R/AA addition)] - [basal respiration].

### Chromatin immunoprecipitation (ChIP)

ChIP-qPCR was performed by adapting the protocol from (Amrute-Nayak et al., 2021). Briefly, Sor- and DMSO-treated myotubes were fixed using ∼1.007% formaldehyde for 10 min at RT, followed by quenching with 125 mM glycine for 5 min. The cells were collected and lysed in ChIP lysis buffer (150 mM NaCl, 50 mM Tris-HCl, pH 7.5, 5 mM EDTA, 0.5 % (v/v) NP-40, 1 % (v/v) Triton X-100, cOmplete EDTA-free Protease inhibitor, PhosSTOP phosphatase inhibitor, 15 mM NEM). The nuclei were collected in Covaris sonication buffer (10 mM Tris-HCl, pH 7.6, 1 mM EDTA, 0.1 % (w/v) SDS). The chromatin was sheared by sonication for 12 min at 10 % df (duty factor), 75 W PIP (peak incident factor) and 200 cycles/burst using the M220 focused-ultrasonicator (#500295, Covaris, US). The chromatin was collected by centrifugation at 16000 g for 15 min at 4°C. 3 % chromatin was collected as input control. The remaining sample was incubated overnight at 4°C with the respective antibodies (2 – 5 µg). IgG was used as the isotype control. Protein G dynabead was added to capture the immunoprecipitated chromatin and bound proteins. After 1.5 h incubation at 4°C, the beads were washed twice with ChIP lysis buffer, thrice with ChIP lysis buffer containing 500 mM NaCl and once with DPBS. The input and samples were heated at 94°C with 10 % (w/v) Chelex-100 to reverse cross-link and purify the DNA. After centrifugation at 16000 g for 1 min at 4°C, the input and ChIPed DNA was collected and processed for qPCR. The percentage of input method was used to analyze the ChIP-qPCR data.

### Total ATP determination

Quantitative estimation of total ATP in C2C12 myotubes was performed using the luciferin-luciferase bioluminescence assay (#A22066, Thermo Scientific, US). Myotubes were collected using trypsin (#12604013, Thermo Scientific, US) and lysed in 300 µl lysis buffer (100mM HEPES pH 7.5, 300 mM NaCl, 1% (v/v) Triton X-100, 2mM EDTA, 2mM EGTA, cOmplete EDTA-free Protease inhibitor, PhosSTOP phosphatase inhibitor) for 20 min at 4°C. After centrifugation at 13000 rpm for 15 min at 4°C, the supernatant was collected and 10 µl sample was added to 90 µl standard reaction solution containing D-luciferin and firefly luciferase. The luminescence was measured using a plate reader at 560 nm. The amount of ATP in the samples was calculated using the standard curve generated from standard ATP solutions. Total protein content (measured using Bradford assay) was used for normalization.

### Mitochondrial membrane potential measurement

Myotubes were stained with 50 nM MitoPT TMRM (tetramethylrhodamine methyl ester) for 30 min, followed by DPBS wash. Positive and negative control of mitochondrial membrane depolarization was generated by treating the myotubes with 50 µM CCCP (carbonyl cyanide 3-chlorophenylhydrazone) and DMSO for 30 min, respectively. Images were acquired using the 20X objective of the Olympus IX83 fluorescent microscope. The accumulation of TMRM was used as a measure of healthy, hyperpolarized mitochondria. To this extent, corrected total cell fluorescence (CTCF) was calculated using the formula-CTCF = Integrated Density – [(Area of selected cell * mean grey value fluorescence of background readings)]

### MitoTracker staining

To visualize the mitochondria, myotubes cultured on laminin-coated coverslips and treated with Sor were incubated with 100 nM MitoTracker Orange (#M7510, Invitrogen, US) for 30 min at 37°C. Following a 15 min wash with DMEM, the cells were fixed using 4 % PFA for 20 min, followed by permeabilization using 0.2 % (v/v) Triton X-100 for 10 min. Standard immunostaining protocol was followed to stain for sarcomeric protein alpha-actinin and the coverslips were mounted on glass slides using fluoroshield (#F6182, Merck, Germany). Images were captured using the 60X oil immersion objective of the Olympus FV1000 confocal microscope (Olympus LifeScience, Japan) at the Research Core Unit for Laser Microscopy, Hannover Medical School, Germany.

### Whole genome transcriptomic analysis (RNASeq)

Four day-differentiated C2C12 myotubes were treated with 20 µM Sor and DMSO for 24 h. Total RNA was isolated using the High Pure RNA Isolation kit (Roche, Switzerland).

#### Library generation, quality control, and quantification

500 ng of total RNA per sample were utilized as input for mRNA enrichment procedure with NEBNext® Poly(A) mRNA Magnetic Isolation Module (#E7490L, New England Biolabs, US) followed by stranded cDNA library generation using NEBNext® Ultra II Directional RNA Library Prep Kit for Illumina (#E7760L, New England Biolabs, US). All steps were performed as recommended in user manual E7760 (Version 1.0_02-2017; NEB), except that all reactions were downscaled to 2/3 of initial volumes.

cDNA libraries were barcoded by dual indexing approach, using NEBNext Multiplex Oligos for Illumina – 96 Unique Dual Index Primer Pairs (#6440S, New England Biolabs, US). All generated cDNA libraries were amplified with 7 cycles of final PCR.

One additional purification step was introduced at the end of the standard procedure, using 1.2x Agencourt® AMPure® XP Beads (#A63881, Beckman Coulter Inc., US). Fragment length distribution of individual libraries was monitored using Bioanalyzer High Sensitivity DNA Assay (#5067-4626, Agilent Technologies, US). Quantification of libraries was performed by use of the Qubit® dsDNA HS Assay Kit (#Q32854, ThermoFisher Scientific, US).

#### Library denaturation and Sequencing run

Equal molar amounts of individually barcoded libraries were pooled for a common sequencing run in which each analyzed library constituted around 13.4 % of overall flowcell/run capacity. The library pool was denatured with NaOH and was finally diluted to 1.8 pM according to the Denature and Dilute Libraries Guide (Document #15048776 v02, Illumina, US). 1.3 ml of the denatured pool was loaded on an Illumina NextSeq 550 sequencer using a High Output Flowcell (400M cluster) for single reads (#20024906, Illumina, US). Sequencing was performed with the following settings-Sequence reads 1 and 2 with 38 bases each; Index reads 1 and 2 with 8 bases each.

#### BCL to FASTQ conversion

BCL files were converted to FASTQ files using bcl2fastq Conversion Software version v2.20.0.422 (Illumina, US).

#### Raw data processing and quality control

Raw data processing was conducted by use of nfcore/rnaseq (version 3.9) which is a bioinformatics best-practice analysis pipeline used for RNA sequencing data at the National Genomics Infrastructure at SciLifeLab Stockholm, Sweden. The pipeline uses Nextflow, a bioinformatics workflow tool. It pre-processes raw data from FastQ inputs, aligns the reads and performs extensive quality control on the results. The genome reference and annotation data were taken from GENCODE.org (*Mus musculus*; GRCm38.p6).

#### Normalization and differential expression analysis

Normalization and differential expression analysis was performed on the internal Galaxy (version 20.05) instance of the RCU Genomics, Hannover Medical School, Germany with DESeq2 (Galaxy Tool Version 2.11.40.6) with default settings except for “Output normalized counts table”, which was set to “Yes” and all additional filters were disabled (“Turn off outliers replacement”, “Turn off outliers filtering”, and “Turn off independent filtering” set “Yes”).

### Quantitative proteomics using mass spectrometry (MS)

#### Sample preparation for MS analysis

Total protein was collected using RIPA buffer (150 mM NaCl, 50 mM Tris-HCl pH 8.0, 1 % Triton X-100, 0.1 % SDS, 0.5 % sodium deoxycholate). Samples at protein concentration of 60 µg were prepared using ROTILoad buffer (Carl Roth, Germany), followed by heating at 95°C for 10 min. The samples were then alkylated by addition of ∼2 % acrylamide, followed by incubation at RT for 30 min. SDS-PAGE was performed on 12 % gels in a mini-protean cell (Bio-Rad Laboratories Inc., US). After electrophoresis, the proteins were stained with Coomassie Brilliant Blue (CBB) for 20 min. Background staining was reduced with water. Each lane was cut into four pieces which were further minced to 1 mm³ gel pieces. Further sample processing was done as described ^62^. Briefly, gel pieces were destained two times with 200 µL 50% ACN, 50 mM ammonium bicarbonate (ABC) at 37°C for 30 min and then dehydrated with 100% ACN. Solvent was removed in a vacuum centrifuge and 100 µl 10 ng/µl sequencing grade Trypsin (Promega, US) in 10 % ACN, 40 mM ABC was added. Gels were rehydrated in trypsin solution for 1 h on ice, and then covered with 10 % ACN, 40 mM ABC. Digestion was performed overnight at 37°C and was stopped by adding 100 µl 50 % ACN, 0.1 % TFA. After incubation at 37°C for 1 h, the solution was transferred into a fresh vial. This step was repeated twice and extracts were combined and dried in a vacuum centrifuge. Dried peptide extracts were redissolved in 30 µl 2 % ACN, 0.1 % TFA with shaking at 800 rpm for 20 min. After centrifugation at 20000 g, aliquots of 12.5 µl each were stored at -20°C.

#### LC-MS analysis

Peptide samples were separated with a nano-flow ultra-high pressure liquid chromatography system (RSLC, Thermo Scientific) equipped with a trapping column (3 µm C18 particle, 2 cm length, 75 µm ID, Acclaim PepMap, Thermo Scientific) and a 50 cm long separation column (2 µm C18 particle, 75 µm ID, Acclaim PepMap, Thermo Scientific). Peptide mixtures were injected, enriched and desalted on the trapping column at a flow rate of 6 µL/min with 0.1% TFA for 5 min. The trapping column was switched online with the separating column and peptides were eluted with a multi-step binary gradient: linear gradient of buffer B (80% ACN, 0.1% formic acid) in buffer A (0.1% formic acid) from 4% to 25% in 30 min, 25% to 50% in 10 min, 50% to 90% in 5 min and 10 min at 90% B. The column was reconditioned to 4% B in 15 min. The Flow rate was 250 nL/min and the column temperature was set to 45°C. The RSLC system was coupled online via a Nano Spray Soure II (Thermo Scientific) to Orbitrap Exploris 240 mass spectrometer. Metal-coated fused-silica emitters (SilicaTip, 10 µm i.d., New Objectives) and a voltage of 2.1 kV were used for the electrospray. Overview scans were acquired at a resolution of 120k in a mass range of m/z 300-1500. Precursor ions of charges two or higher and a minimum intensity of 4000 counts were selected for HCD fragmentation with a normalized collision energy of 38.0, an activation time of 10 ms and an activation Q of 0.250. Active exclusion was set to 70 s within a mass window of 10 ppm of the specific m/z value.

Raw MS data were processed using Max Quant software, version 2.0 ^62^ and Perseus software, version 2.0.6.0, ^63^ and mouse entries of uniprot DB. Proteins were stated identified by a false discovery rate of 0.01 on protein and peptide level.

### Myotubes length and diameter measurement

Myotube length and thickness was measured for control and drug treated cells using ImageJ analysis tools. As the diameter of the Sor-treated myotubes varied along the length of the cell, the myotube was checked approximately every 1 µm apart to accurately estimate the diameter comprising different sections. Highly varied values were noted even within single cells in contrast to the control cells that remained relatively uniform throughout. Note that the average myotube lengths presented in the plots for control, Ima, and Nilo are underestimated, as a large fraction of myotubes extend beyond even the lower magnified field of view and thus could not be measured. Therefore, the difference with the Sor-treated cells could be even larger.

### Kymograph

The distance over time plot reveals the myotubes contractions. ImageJ plug-in ‘multiple kymograph’ was used to measure the movement of intracellular structures in response to electrical stimulation of the myotubes as the cell underwent contraction-relaxation cycles. Representative kymographs corresponding to short sections of intra-myotubes components are compared when cells were treated with different drugs.

## Acknowledgements

We thank Dr. Sarah Elsheikh and Prof. Dr. Susanne Häußler (Institute for Molecular Bacteriology, TWINCORE GmbH, Center of Clinical and Experimental Infection Research, A Joint Venture of the Hannover Medical School and the Helmholtz Center for Infection Research, Hannover, Germany) for the Seahorse XFe96 analyzer. We thank PD Dr. Ortwin Naujok (Institute for Clinical Biochemistry, Hannover Medical School, Germany) for the microplate reader. RNAseq data used in this publication was generated by the Research Core Unit Genomics (RCUG) at Hannover Medical School, Germany. We acknowledge the support of Dr. Chiara Cordiglieri and Dr. Alessandra Fasciani from the Imaging Unit of Istituto Nazionale Genetica Molecolare (INGM) (Marco Ghilotti).

## Authors contributions

AN and MA conceived the project. AN and BK designed the experiments. BK performed most of the experiments analysed and interpreted the data, and wrote the manuscript. PS, VR, and CL isolated satellite stem cells from mice, prepared primary myotubes, prepared RNA and cDNA, and performed immunostaining and microscopy. AP performed the mass spectrometry. TK provided her critical expertise in muscle physiology. MA contributed to designing the experiments, data analysis, interpretation of the data, and editing the manuscript. AN supervised the project, analysed and interpreted the results, wrote and edited the manuscript.

## Funding

This research was partly supported by grants from Deutsche Forschungsgemeinschaft (DFG) to AN (NA 1565/2-1), Deutsche Krebshilfe to AN (refernce number: 70115510) and a grant from Fritz Thyssen Stiftung to MA (10.19.1.009MN).

## Ethics approval

All the experimental procedures were performed under the ethical approval of the Italian Ministry of Health and the Institutional Animal Care and Use Committee (authorization no. 83/2019-PR). The animals were maintained in an authorized facility at San Raffaele Hospital, Milan (authorization no. N. 127/2012-A).

## Statistical analysis

All statistical analysis was performed using GraphPad Prism 9.5.1. Unpaired t-test with Welch’s correction was used to determine the statistical significance in terms of p-value.

## Declaration

The author declare no competing interests.

## Supplemental Information

The supplemental information contains Fig. S1 – Fig.S5

**Supplementary Fig. 1:**
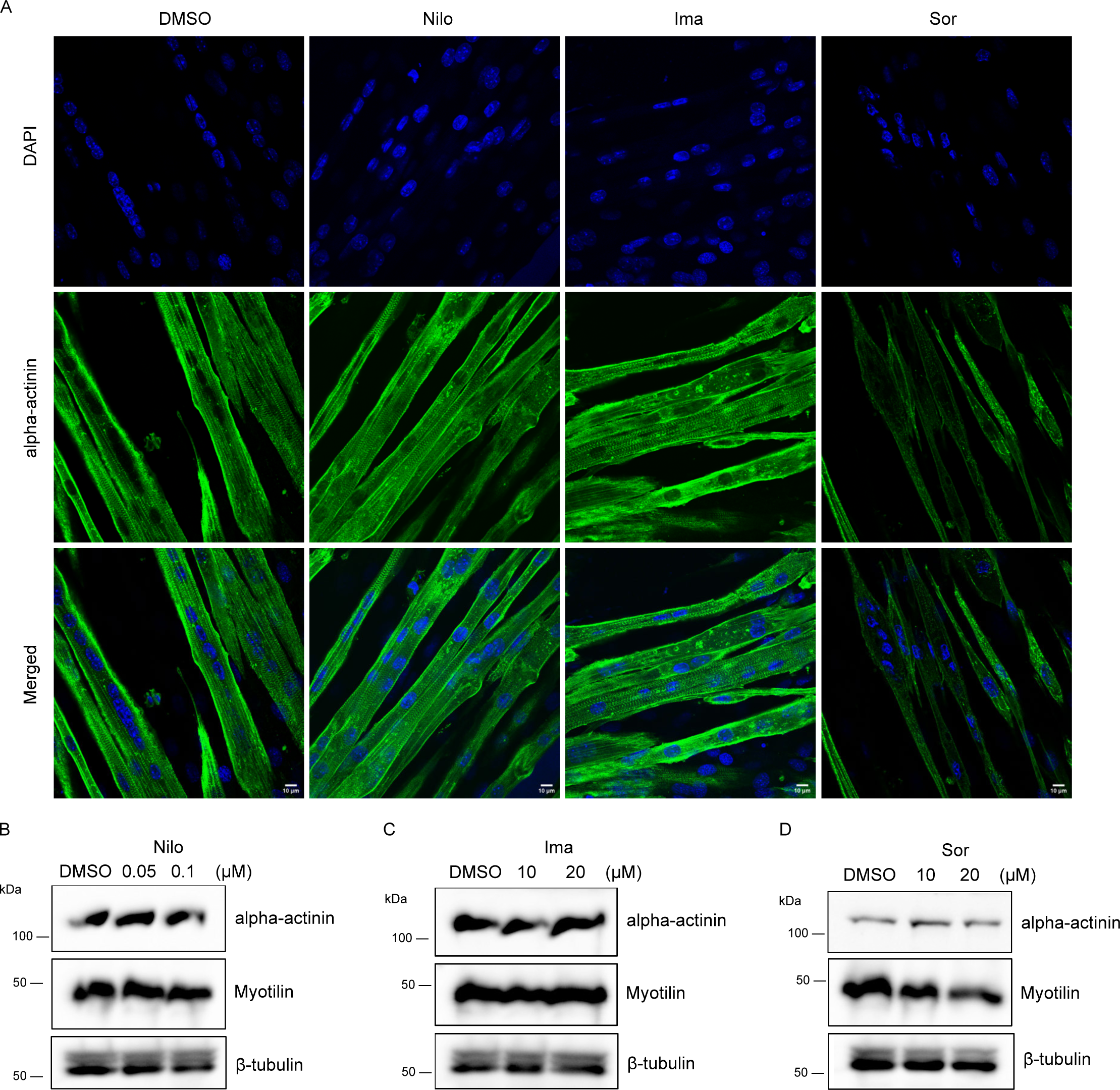
TKI treatment of C2C12 myotubes. (A) Representative images of TKI-treated myotubes immunostained against alpha-actinin (green) and nucleus (DAPI – blue). Images were captured using the 60X oil immersion objective. Scale bar = 10 µm. (B-D) Western blot shows protein expression of alpha-actinin and myotilin in (B) Nilo, (C) Ima, and (D) Sor treated myotubes, along with the respective DMSO control. β-tubulin was used as the loading control. All concentrations are in µM.

**Supplementary Fig. 2:**
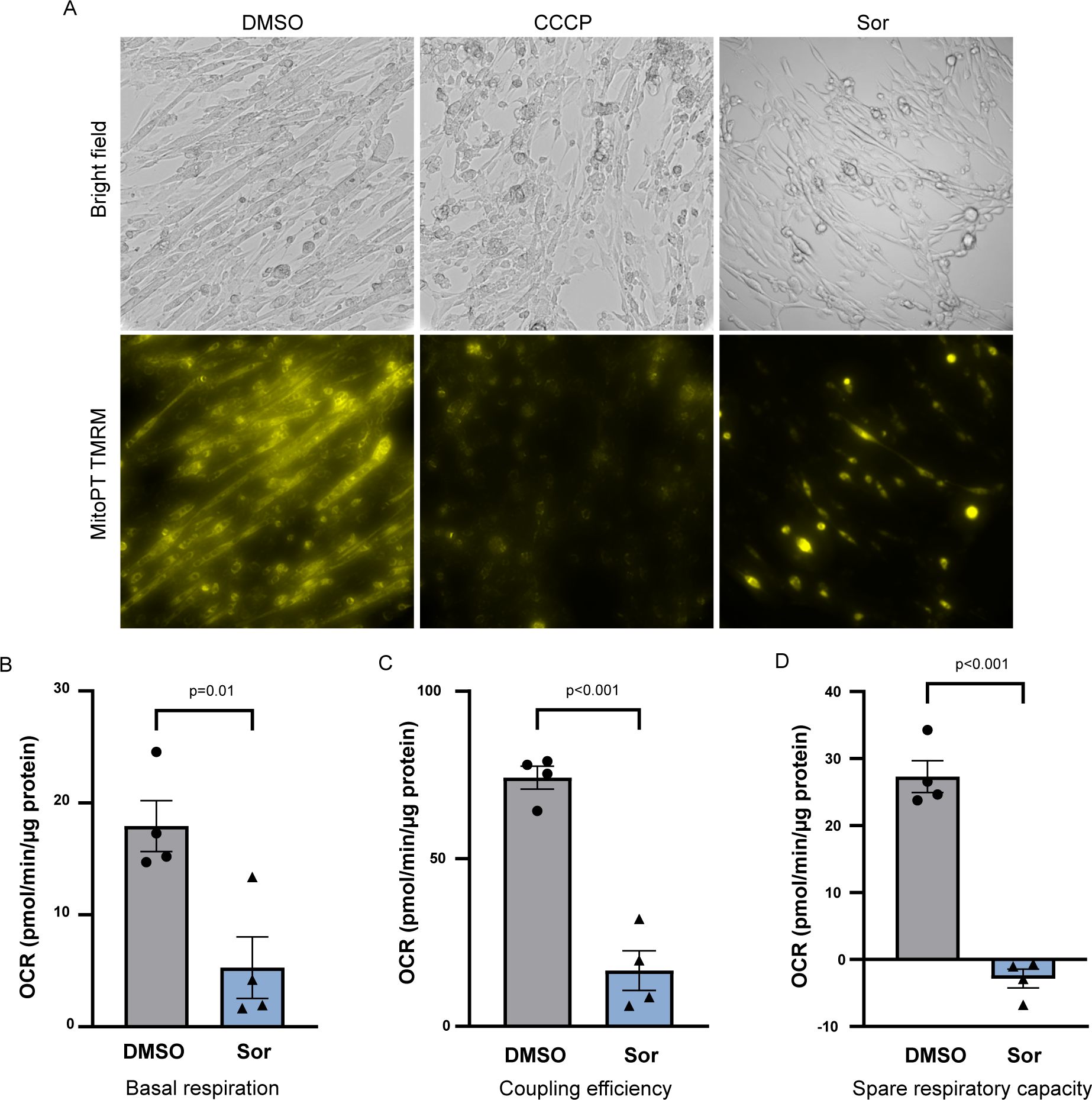
Sorafenib treatment leads to mitochondrial dysfunction in muscle cells. (A) Representative images of Sor-treated myotubes stained with MitoPT TMRM dye (yellow). Images were captured using the 20X objective. CCCP was used as the positive control of mitochondrial membrane depolarization. (B-D) Graph shows (B) basal respiration, (C) coupling efficiency, and (D) spare respiratory capacity, of DMSO- and Sor-treated myotubes, as calculated from the OCR readings obtained in the Seahorse MitoStress Test from Fig. 3F. (Mean ± SEM, N = 4, n = 44). Statistical significance was determined using unpaired t-test with Welch’s correction on GraphPad Prism 9.5.1. DMSO = grey, Sor = blue.

**Supplementary Fig. 3:**
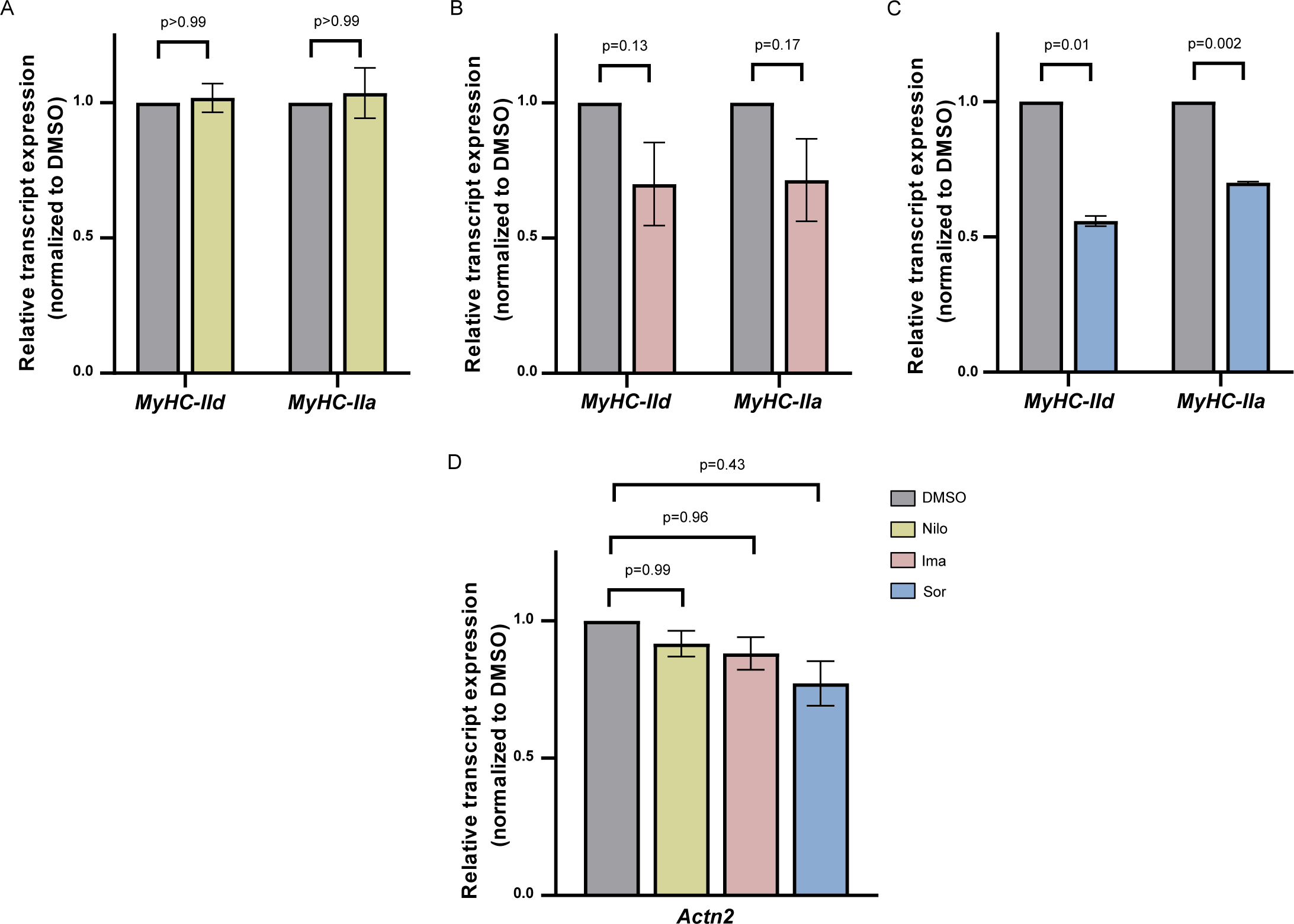
Effect of TKI treatment on transcript expression of sarcomeric genes in myotubes. Graph shows relative transcript expression of MyHC-IId and –IIa in myotubes treated with (A) Nilo, (B) Ima, and (C) Sor. (D) Graph shows relative transcript expression of Actn2 in myotubes treated with TKIs. The transcript expression for each gene was normalized to Gapdh and its respective DMSO control (grey). All graphs show Mean ± SEM, where N = 3, n = 9. Statistical significance was determined using unpaired t-test with Welch’s correction on GraphPad Prism 9.5.1. The concentrations used for Nilo was 0.1 μM, and 20 μM for Ima and Sor. DMSO = grey, Nilo = green, Ima = red, Sor = blue.

**Supplementary Fig. 4:**
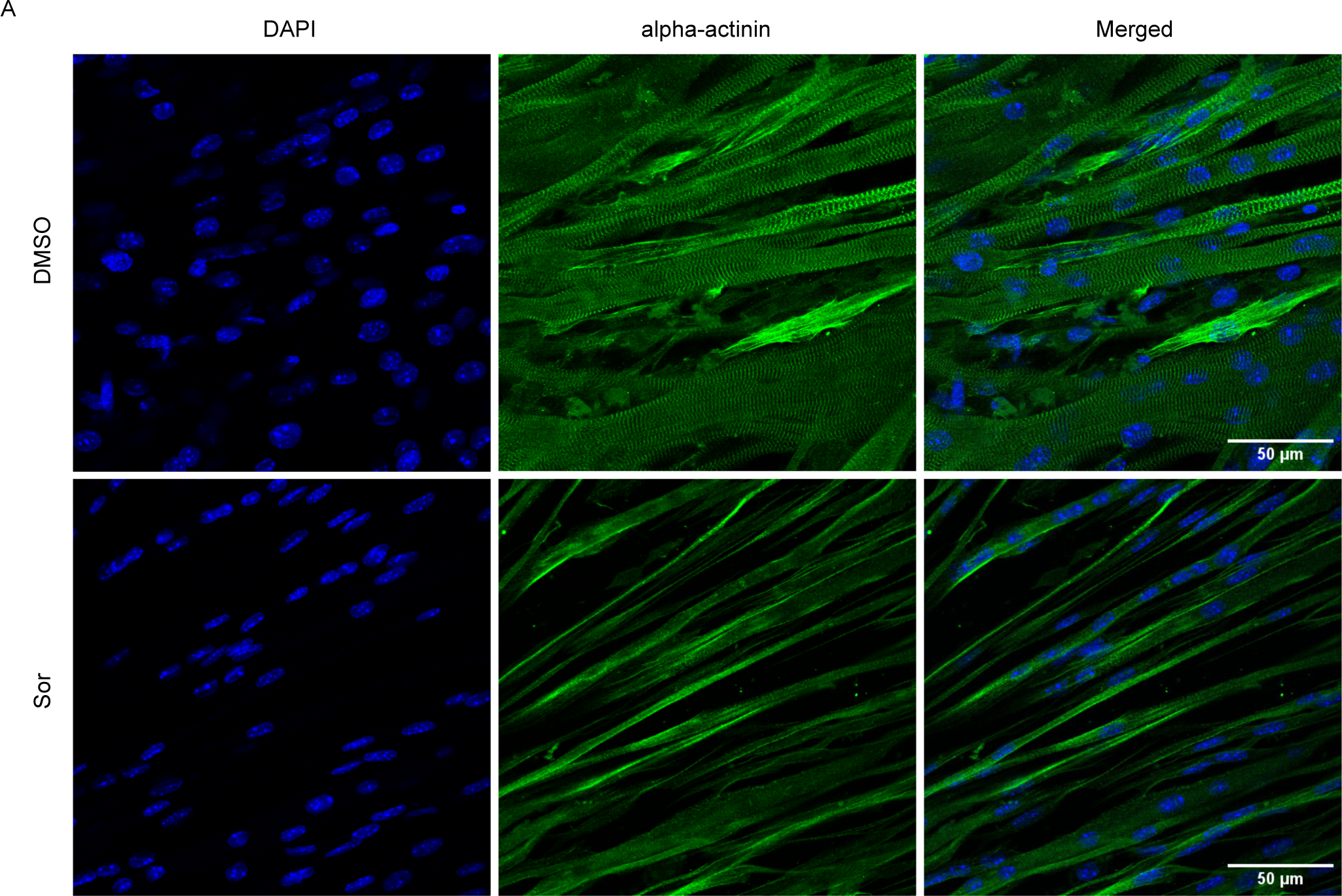
Sorafenib disrupts sarcomere organization in satellite cell-derived muscle cells. (A) Representative images of satellite cell-derived muscle cells immunostained against alpha-actinin (green) and nucleus (DAPI – blue). Scale bar = 50 μm.

**Supplementary Fig. 5:**
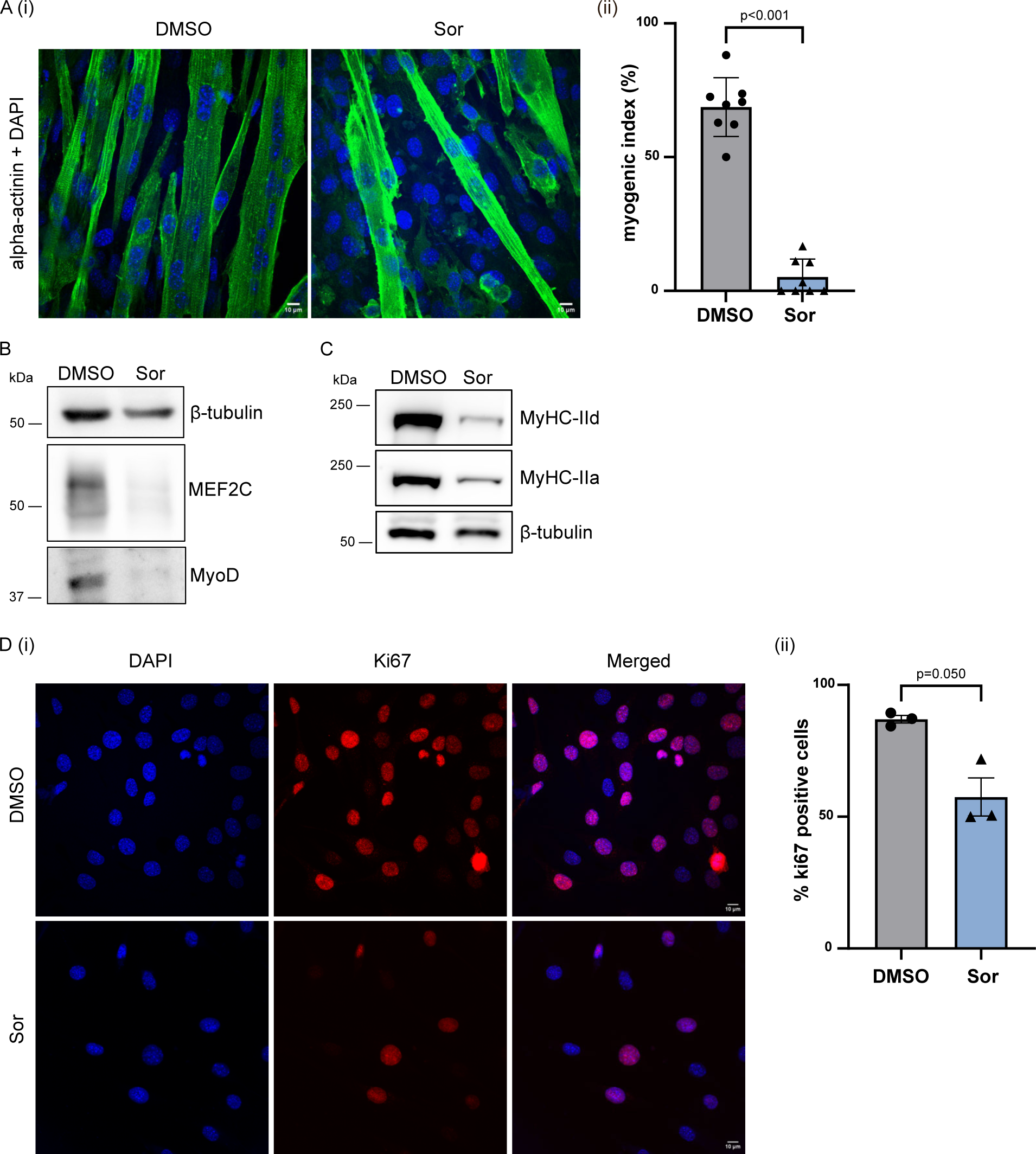
Sorafenib impairs myogenic differentiation process. (A.i) Representative overlaid images of myotubes generated from DMSO- and Sor-treated myoblasts, immunostained against alpha-actinin (green) and nucleus (DAPI – blue). Scale bar = 10 µm. (A.ii) Graph shows myogenic index (%) as calculated from (A.i) (Mean ± SEM, DMSO – n = 407, Sor – n = 416, where n = total number of cells). Western blot shows protein expression of (B) myogenic regulatory factors – MEF2C and MyoD, (C) motor proteins – MyHC-IId and -IIa, in myotubes generated from DMSO- and Sor-treated myoblasts. β-tubulin was used as the loading control. (D.i) Representative confocal microscopy images of DMSO- and Sor-treated myoblasts immunostained against proliferation marker Ki67 (red) and nucleus (DAPI – blue). Scale bar = 10 µm. (D.ii) Graph shows the percentage of Ki67-positive cells, as counted from (D.i) (Mean ± SEM, N = 3, DMSO – n = 680, Sor – n = 389, where N = biological replicates and n = total number of nuclei). Statistical significance was determined using unpaired t-test with Welch’s correction on GraphPad Prism 9.5.1. DMSO = grey, Sor = blue.

**Supplementary movie 1: Contractile ability of Nilo-treated myotubes in comparison to control (DMSO) cells.** The rhythmic contraction (excitation-contraction coupling) of DMSO (left frame) and Nilo-treated (right frame) myotubes on laminin-coated coverslips, in response to electrical stimulation at 40 V and 1 Hz, was recorded at 10 fps. The movies are played at 20 fps. Scale bar of 200 pixel corresponds to 58 µm. The myotubes are pseudo-coloured.

**Supplementary movie 2: Contractile ability of Ima-treated myotubes compared to control (DMSO) cells.** The movies are played at 20 fps. Scale bar of 200 pixel corresponds to 58 µm. DMSO treatment is on the left frame, Ima-treated is on the right.

**Supplementary movie 3: Contractile ability Sor-treated myotubes in comparison to control (DMSO) cells.** The movies are played at 20 fps. Scale bar of 200 pixel corresponds to 58 µm. DMSO treatment is on the left frame, Sor-treated is on the right.

**Supplementary Table 1:**

Data from RNASeq assays (supplementary table 2) and proteomic analysis (supplementary table 3) was used for DAVID GO analyses to annotate the significantly enriched GO terms among the two datasets.

**Sheet 1,** List of genes/proteins associated with the specific GO terms that were differentially regulated (up- or down-regulated) in our transcriptomic dataset (labelled as ‘RNASeq_regulated’ and proteomic dataset (labelled as ‘MS_regulated’. The highlighted genes are unique to that specific dataset (RNASeq or MS), i.e, these candidates showed regulated response only either in RNAseq or MS. GO terms associated with Myofibril include the following GO terms: contractile fiber, myofibril, myofibril assembly, sarcomere organization, muscle contraction. GO terms associated with SR include the following GO terms: sarcoplasmic reticulum, calcium ion binding. GO terms associated with Mitochondria include the following GO terms: mitochondria, mitochondrial membrane, mitochondrial inner membrane, mitochondrial envelope, mitochondrial respirasome, mitochondrial calcium ion transmembrane transport, oxidative phosphorylation, glycogen metabolic process.

**Sheet 2,** List of genes/proteins associated with the specific GO terms, that were identified to be downregulated in our proteomic dataset (labelled as ’MS_down’), but remained unchanged in the transcriptomic dataset (labelled as ’RNASeq_unchanged’). GO terms associated with Myofibril include the following GO terms: contractile fiber, myofibril, myofibril assembly, sarcomere organization, muscle contraction. GO terms associated with SR include the following GO terms: sarcoplasmic reticulum, calcium ion binding. GO terms associated with Mitochondria include the following GO terms: mitochondria, mitochondrial membrane, mitochondrial inner membrane, mitochondrial envelope, mitochondrial respirasome, mitochondrial calcium ion transmembrane transport, oxidative phosphorylation, glycogen metabolic process.

**Sheet 3,** List of genes/proteins associated with the specific GO terms, that were identified to be differentially regulated in our proteomic dataset (labelled as ’MS_reg’) and the transcriptomic dataset (labelled as ’RNASeq_reg’). Upregulated genes/proteins are indicated in orange, while downregulated ones are indicated in blue. GO terms associated with ER include the following GO terms: endoplasmic reticulum quality control compartment, endoplasmic reticulum chaperone complex, smooth endoplasmic reticulum, endoplasmic reticulum unfolded protein response, response to unfolded protein, response to endoplasmic reticulum stress, endoplasmic reticulum lumen, endoplasmic reticulum membrane, endoplasmic reticulum.

**Supplementary Table 2:**

Differential gene expression analysis (RNASeq assays) was performed using the DESeq2 tool (Galaxy Tool Version 2.11.40.6). The original file contains list of all identified genes, their log2fold-change and p-value in DMSO vs Sor treatment.

**Supplementary Table 3:**

Raw MS data was processed using MaxQuant v2.0 and Perseus v2.0.6.0 to identify changes in global proteome upon Sor treatment. The original file contains list of all identified proteins, their log2fold-change and –log10(p-value) in DMSO vs Sor treatment.

## Notes

### Competing Interest Statement

The authors have declared no competing interest.

